# A novel batokine Breg controls adipose thermogenesis and glucose homeostasis

**DOI:** 10.1101/2022.08.16.504121

**Authors:** Qingbo Chen, Lei Huang, Dongning Pan, Kai Hu, Rui Li, Lihua J. Zhu, David A. Guertin, Yong-Xu Wang

## Abstract

Batokines selectively expressed in brown and beige adipocytes remain to be identified and their potential signaling role in adipose thermogenesis are largely unknown. Here we identified a batokine we named as Breg acting as a key regulator for adipose thermogenesis and glucose homeostasis. Breg expression is adipose-specific and highly brown fat-enriched, and its secretion is stimulated by β3-adrenergic activation. Gain-of-functional studies collectively showed that secreted Breg promotes adipose thermogenesis, lowers glucose level, and protects against obesity. Adipose-specific Breg knockout mice are defective in white fat browning, and are susceptible to high fat diet-induced obesity and hyperglycemia, demonstrating the physiological importance of this batokine in energy metabolism. Mechanistically, Breg binds to a putative receptor on adipocyte surface and activates protein kinase A independently of β-adrenergic signaling. These results establish Breg as a major upstream signaling component in thermogenesis and offer a potential avenue for the treatment of obesity and diabetes.

---

White fat (WAT) browning and non-shivering thermogenesis are principally controlled by β-adrenergic receptor (β-AR) signaling^1, 2^. Studies in mice have demonstrated that brown fat (BAT) and beige adipocytes can not only protect against obesity but also improve glucose homeostasis independent of changes of body weight^3-8^, although the underlying mechanisms of the latter effect remain unclear^8^. While agonists of β3-adrenergic receptor (β3-AR) have exhibited efficacy in energy expenditure and glucose uptake in human studies, their undesirable cardiovascular effects prevent their clinical use^9, 10^. Therefore, there is an urgent need to identify thermogenic activators or signaling pathways that act independently of β-AR signaling.

While adipokines secreted from WAT have been extensively investigated, the secretomes of BAT and beige adipocytes have not been well characterized, let alone functionally studied. Although a few brown adipokines (batokines) have been identified, they are highly expressed in other tissues and/or have a very low level in BAT (and WAT)^11, 12^. Moreover, their physiological functions attributable to adipose tissue secretion remain to be explored^12^. Thus, the roles of BAT- and beige-selective batokines in energy homeostasis are largely unknown.

## Results

### Breg is a novel adipokine highly enriched in BAT and beige adipocytes and its secretion is stimulated by β3-adrenergic activation

We aimed to identify novel BAT- and beige-selective batokines that can act on adipose tissue to induce WAT browning. After removal of small molecules, serum-free conditioned medium collected from differentiated brown adipocyte culture was able to stimulate the expression of BAT-selective genes including *Ucp1* in primary inguinal adipocytes (Extended Data Fig. 1a), whereas heat-inactivated medium had little effect, suggesting that brown adipocytes secrete a protein(s) with browning activity. We exploited three independent strategies to identify this potential secreted protein (Fig. 1a). We first screened our previously published RNA-Seq datasets^13^ and identified 139 genes that are both abundantly and selectively expressed in BAT relative to epididymal WAT and skeletal muscle. We then performed proteomic analysis of serum-free conditioned medium collected from differentiated brown adipocytes and identified 1894 proteins. Among them, 1751 protein were also identified in conditioned medium of iWAT adipocytes^14^. Combining our proteomics data with the RNA-seq data led to 42 candidates (Extended Data Table 1). Interestingly, many of them are mitochondrial proteins. We envisioned that these mitochondrial proteins are likely secreted through extracellular vesicles, as mitochondrial components constitute a major part of extracellular vesicles^15^ and brown fat-derived exosomes^16^. Lastly, we used bioinformatics to predict whether any of the 42 candidates are potentially secreted proteins, and identified two proteins, Apolipoprotein C-I (Apoc1) and UPF0687. We focused on UPF0687, which is encoded by *1700037H04Rik. 1700037H04Rik* and its human orthologue *C20orf27* are previously uncharacterized genes, and encode 19 kilodalton (kDa) products (Extended Data Fig. 1b) that are predicted by SecretomeP^17^ to be non-classically secreted proteins. Five peptides of this gene product were identified in our proteomic analysis (Fig. 1b). We named this adipokine as Breg (<u>B</u>rowning <u>reg</u>ulator).

**Fig. 1.**
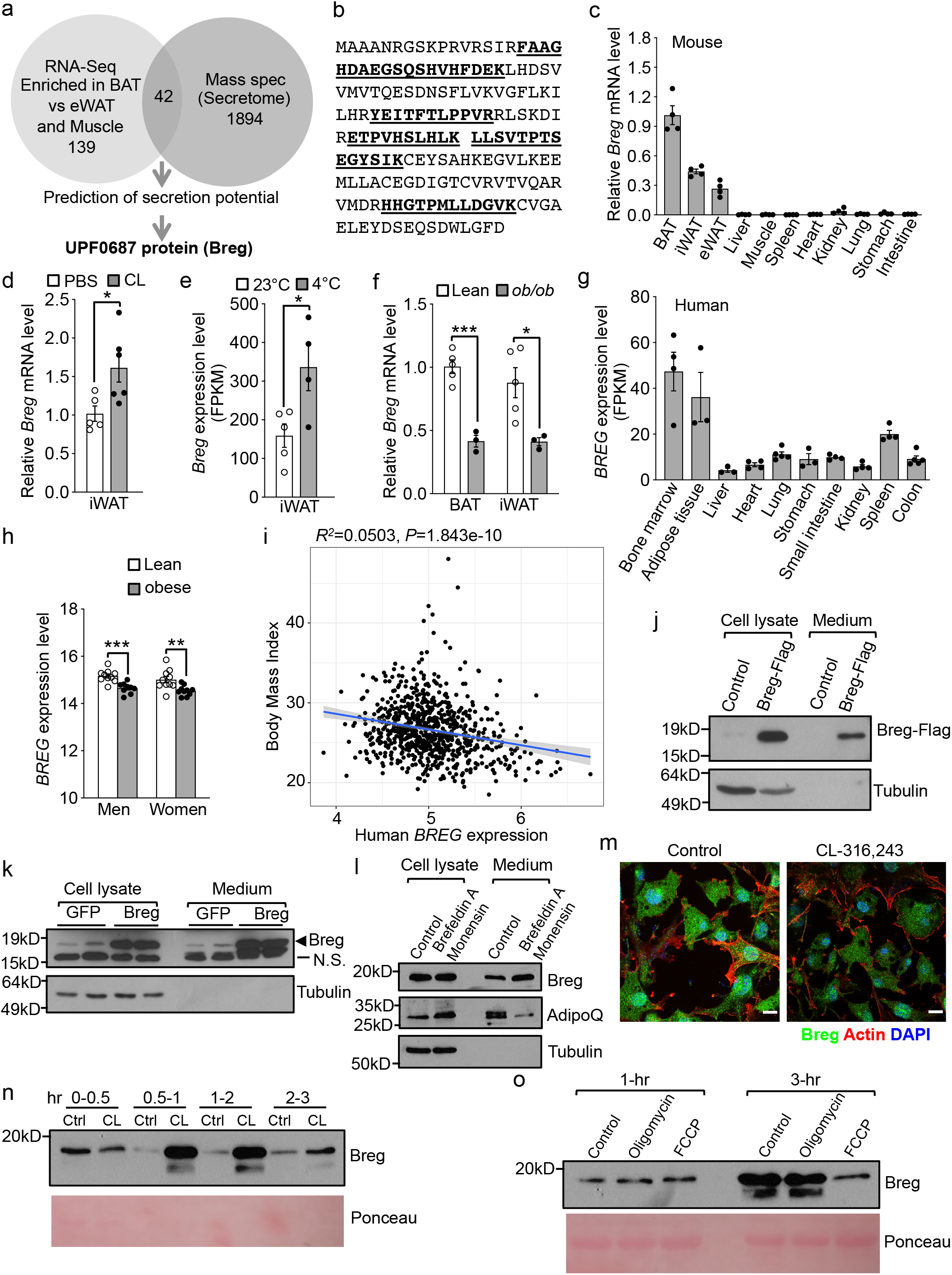
Identification of Breg. **a**, Strategies to identify brown adipocyte secreted proteins. **b**, Breg matched peptides identified by mass spectrometry are bold and underlined. **c**, *Breg* expression in wild type (WT) mice tissues (n=4). **d**, *Breg* expression in inguinal WAT (iWAT) from WT female mice administrated with PBS (n=5) or CL-316,243 (CL) (n=6). **e**, *Breg* expression in iWAT from WT mice housed at 22 °C (n=5) or cold (4 °C) (n=4) for 3 days from published data^18^. **f**, *Breg* mRNA expression in BAT and iWAT from male *ob/ob* mice and lean controls (n=3-5). **g**, Human *BREG* expression in different tissues (n=3-5) from published data^19^. **h**, Expression of human *BREG* in subcutaneous adipocytes from lean and obese subjects (n=9-10) from published data^20^. **i**, Linear regression analysis of BMI and *BREG* expression from adipose tissue of 770 men from published data^21^. **j**, Detection of secreted Breg-Flag from brown adipocytes containing a Flag knock-in. **k**, Detection of Breg secretion from brown adipocytes. N.S., non-specific band. **l**, Detection of Breg secretion from brown adipocytes treated with Brefeldin A and Monensin for 6 hours. **m**, Subcellular localization of Breg in brown adipocytes treated with or without CL-316,243 for 45 minutes. Scale bar, 20µm. **n**, Detection of Breg secretion from brown adipocytes treated with CL-316,243 at indicated time intervals. **o**, Detection of Breg secretion from brown adipocytes treated with oligomycin or FCCP for 1 hour or 3 hours. Data are mean ± s.e.m. *p<0.05, **p<0.01, ***p<0.001 by two-tailed Student’s t test.

Quantitative PCR (qPCR) analysis showed that *Breg* expression was restricted to adipose tissue and highly enriched in BAT, and was primarily present in mature adipocytes (Fig. 1c and Extended Data Fig. 1c, d). Daily β3-AR agonist CL-316,243 administration for 2 days and cold challenging at 4°C for 3 days led to increased expression of *Breg* in inguinal WAT (Fig. 1d, e)^18^, while acute treatment of adipocyte culture with CL-316,243 in vitro or acute cold exposure in vivo had no effect (Extended Data Fig. 1e, f). *Breg* expression was decreased in BAT and inguinal WAT of *ob/ob* mice (Fig. 1f). We also analyzed publicly available gene expression datasets, and found that *BREG* expression was enriched in human adipose tissue (Fig. 1g)^19^, and was lower in adipocytes of obese human subjects compared with those of lean subjects (Fig. 1h)^20^. Importantly, we performed linear regression analyses on microarray data from subcutaneous adipose tissue samples of a cohort of 770 men and found that levels of *BREG* expression were negatively correlated with body mass index (BMI) (Fig. 1i)^21^.

To further confirm Breg is a secreted protein, we used CRISPR/Cas9 system to knock-in a Flag tag at endogenous Breg locus immediately before its stop codon in immortalized brown preadipocytes. When probed with a Flag antibody, Breg-Flag was detected in conditioned medium of differentiated knock-in adipocytes (Fig. 1j). We then obtained a commercial Breg antibody that was generated against human BREG. Western blot analysis of BAT extracts from *Breg* knockout mice validated the antibody (Extended Data Fig. 1g). Despite that the antibody had a relatively low reactivity to mouse Breg protein (Extended Data Fig. 1h), we were able to detect endogenous Breg present in conditioned medium of mouse brown adipocyte culture, and its level was increased when brown adipocytes were infected with adenovirus expressing *Breg* (Fig. 1k). Based on equivalent loading of cell numbers, we estimated that at least 50% of produced Breg in adipocyte culture was secreted into medium during a 16-hr incubation. Similarly, in a competitive ELISA assay we developed (Extended Data Fig. 1i), we found that about 10% of total Breg protein was secreted during a 3-hr incubation. Moreover, BREG was shortlisted as a secreted protein in proteomic analyses of human adipocyte secretome^22^. Together, these results confirmed our proteomic data that endogenous Breg is an adipokine.

Brefeldin A and monensin together block classical protein secretory pathway from the ER to the trans-Golgi apparatus cisternae. While adiponectin secretion was inhibited by brefeldin A and monensin as expected, Breg secretion was instead increased (Fig. 1l). It has been reported that brefeldin A and monensin enhance the secretion of non-classically secreted proteins IL-1β and migration inhibitory factor^23, 24^. These data provide additional evidence that Breg is secreted via a non-classical secretory pathway.

We found that Breg was present with a punctate or vesicular pattern (Fig. 1m). Importantly, treatment of adipocytes with β3-AR agonist CL-316,243 decreased Breg staining (Fig. 1m), raising the possibility of increased secretion. To investigate this further, we collected conditioned medium at different time intervals. We found that Breg secretion was stimulated by CL-316,243 during the first 2-hr treatment, and this stimulation waned afterwards (Fig. 1n). Thus, β3-AR activation causes a spike of Breg secretion.

Interestingly, FCCP, an uncoupler of mitochondrial oxidative phosphorylation, inhibited Breg secretion, whereas oligomycin, an inhibitor of ATP synthase, had no effect (Fig. 1o), which suggests that the uncoupling process, not ATP level per se, feeds back to negatively regulate Breg secretion. The dynamic regulation of Breg secretion by β3-AR activation and FCCP supports a potential role of this batokine in adipose thermogenesis.

### Transgenic expression of *Breg* enhances energy expenditure, improves glucose homeostasis and protects against diet-induced obesity

Acute overexpression of *Breg* with adenovirus in primary inguinal adipocytes increased expression of BAT markers *Ucp1, Cidea* and genes involved in mitochondrial oxidative metabolism and glycolysis (Extended Data Fig. 2a). General adipogenesis was not affected (Extended Data Fig. 2a, b). Co-treatment with CL-316,243 had an additive effect on *Ucp1* level (Extended Data Fig. 2c). Moreover, conditioned medium from HEK293 cells overexpressing *Breg* induced *Ucp1* expression in primary inguinal adipocytes (Extended Data Fig. 2d). To determine whether Breg is capable of driving WAT browning in vivo, we generated *aP2-Breg* transgenic mice that selectively overexpressed *Breg* in adipose tissue (Extended Data Fig. 3a). Serum collected from *Breg* transgenic mice, which had a higher Breg level in circulation (Extended Data Fig. 3b), increased thermogenic genes expression in inguinal adipocyte culture (Extended Data Fig. 3c). On a normal chow diet, while there was no difference in body weight and food intake (Extended Data Fig. 3d, e), the transgenic mice had elevated expression of thermogenic and glycolytic genes in inguinal WAT, but not in BAT or epididymal WAT, compared with littermate controls (Fig. 2a, b and Extended Data Fig. 3f, g). Haemotoxylin and Eosin (H&E) staining and Ucp1 immunofluorescence staining confirmed robust inguinal WAT browning that possessed numerous small, UCP1-positive adipocytes (Fig. 2c and Extended Data Fig. 3h). Consistent with this, injection of Breg adenovirus into the inguinal WAT also promoted its browning (Extended Data Fig. 3i, j). As expected, the transgenic mice displayed a significantly higher oxygen consumption rate in inguinal WAT (Fig. 2d), and their body temperature is higher compared with that of control mice at cold (Fig. 2e). These data revealed that Breg promotes thermogenic activation and augments energy expenditure.

**Fig. 2.**
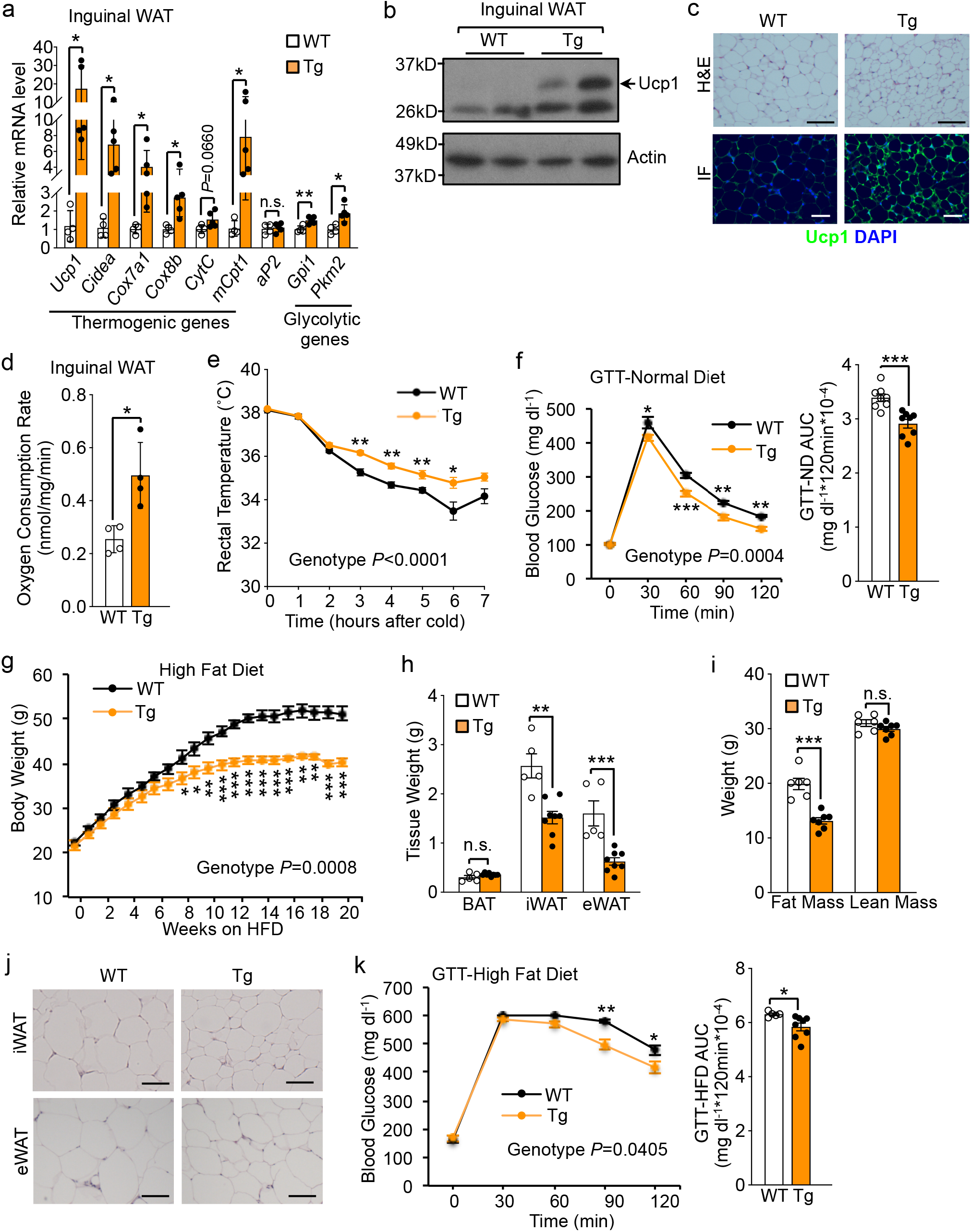
Targeted expression of *Breg* enhances energy expenditure. **a**, Gene expression in iWAT from 2-month-old male *Breg* transgenic (Tg) mice (n=5) and littermate controls (n=4). **b**, Western blot of Ucp1 in iWAT from *Breg* Tg mice. **c**, Representative images of H&E staining and Ucp1 immunofluorescence (IF) staining in iWAT from *Breg* Tg mice (n=3 mice per group). Scale bar, 200 µm. **d**, Oxygen consumption rate in iWAT from *Breg* Tg mice (n = 4 mice per group). **e**, Rectal temperature of 4-month-old female *Breg* Tg mice (n=9) and littermate controls (n=5) during cold exposure. **f**, Glucose tolerant test (GTT) in 10-month-old male *Breg* Tg mice and littermate controls on normal chow diet (n=8 mice per group). **g**, Body weight of male *Breg* Tg mice (n=8) and littermate controls (n=5) on high-fat diet (HFD). **h**, Fat mass of mice from (**g**) after 24 weeks of HFD feeding. **i**, Fat mass and lean mass of a second cohort of mice after 16 weeks of HFD feeding (WT, n=6; Tg, n=7). **j**, Representative images of H&E staining of iWAT and eWAT from mice in (**g**) after 24 weeks of HFD feeding. Scale bar, 200 µm. **k**, GTT in mice from (**g**) on 16 weeks of HFD. Data are mean ± s.e.m. *p<0.05, **p<0.01, ***p<0.001 and not significant (n.s.) by two-tailed Student’s t test (**a, d, h, i** and AUC in **f** and **k**) and two-way repeated measures ANOVA with post hoc test by Fisher’s LSD test (**e, f, g** and **k**).

The *Breg* transgenic mice were more glucose tolerant (Fig. 2f), consistent with the notion that WAT browning improves glucose homeostasis^3, 5, 6, 25-27^. On a high-fat diet (HFD), the transgenic mice gained significantly less body weight and fat mass (Fig. 2g, h), despite similar food intake as control mice (Extended Data Fig. 3k). This was confirmed in a second cohort of HFD-fed mice (Fig. 2i and Extended Data Fig. 3l). Inguinal WAT and epididymal WAT adipocytes are substantially smaller (Fig. 2j). Moreover, the transgenic mice fed on HFD also displayed better glucose homeostasis (Fig. 2k). Together, inguinal WAT browning in the *Breg* transgenic mice improves glucose homeostasis and protects against HFD-induced obesity.

### Breg is required for inguinal WAT browning

To investigate the thermogenic function of endogenous Breg, we generated *Breg flox/flox* (Flox) mice, and then crossed the Flox mice with *Adiponectin-Cre* mice^28^ to generate adipose-specific *Breg* knockout (ADKO) mice. Analysis of *Breg* mRNA expression from multiple tissues confirmed that the deletion was specific to WAT and BAT (Extended Data Fig. 4a). At room temperature with a normal chow diet, the ADKO mice had similar body weight and food intake as littermate control mice, and no difference in thermogenic genes expression was observed in adipose tissues including inguinal WAT (Extended Data Fig. 4b-4f). This prompted us to ask whether Breg is required for beige fat biogenesis at thermogenic demanding conditions. As expected, we found that cold exposure stimulated the generation of UCP1-positive beige adipocytes in the inguinal WAT of control mice; however, beige fat formation was strikingly impaired in ADKO mice (Fig. 3a), which was further confirmed by analysis of *Ucp1* mRNA expression (Fig. 3b). Correspondingly, the body temperature was lower in the ADKO mice than control mice (Fig. 3c). Interestingly, ADKO mice contained more inguinal WAT mass after acute cold exposure (Extended Data Fig. 4g), reflecting decreased fat burning. Compromised inguinal WAT browning also led to decreased glucose uptake (Fig. 3d). We next determined whether Breg is indispensable for β3-adrenergic signaling-induced inguinal WAT browning. Similar to what occurred at condition of cold challenging, administration of CL-316,243 led to robust inguinal WAT browning in control mice but not in ADKO mice (Fig. 3e and Extended Data Fig. 4h), and ADKO lost less inguinal WAT mass (Extended Data Fig. 4i). Together, these results suggest that endogenous Breg is critically required for both cold- and β3-adrenergic agonist-stimulated beige fat formation and thermogenesis.

**Fig. 3.**
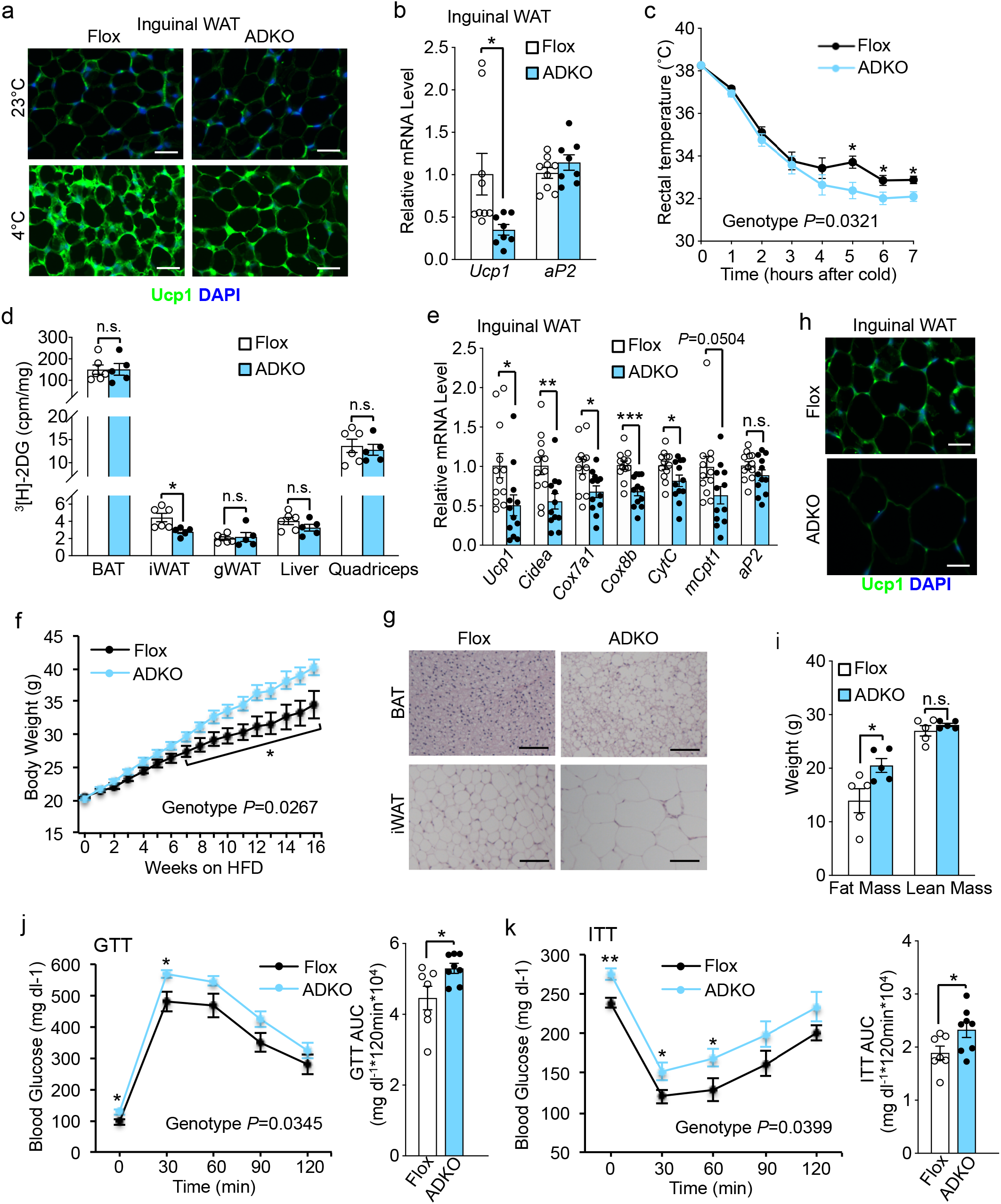
Breg is required for iWAT browning and energy homeostasis. **a**, Representative images of Ucp1 immunofluorescence staining in iWAT from 3-month-old male *Breg* ADKO mice and Flox controls (n=3 mice per group). Scale bar, 200 µm. **b**, Gene expression analysis in iWAT from 5-month-old female *Breg* ADKO mice (n=8) and Flox controls (n=9) after 7 hours cold exposure. **c**, Rectal temperature of mice from (**b**) during cold exposure. **d**, Quantification of ^3^H-2DG uptake in indicated tissues from *Breg* ADKO mice (n=5) and Flox controls (n=6) after 3 hours cold exposure. **e**, Gene expression analysis in iWAT from CL-316,243 administrated mice (n=12 mice per group). **f**, Body weight of male *Breg* ADKO mice (n=15) and Flox controls (n=12) on HFD. **g, h**, Representative images of H&E staining in BAT and iWAT (**g**) and Ucp1 immunofluorescence staining in iWAT (**h**) from mice in (**f**) on 21 weeks of HFD (n=3 mice per group). Scale bar, 200 µm. **i**, Fat mass and lean mass of a second cohort of mice after 16 weeks of HFD feeding (n=5 mice per group). **j**, GTT in mice from (**f**) on 18 weeks of HFD (Flox, n=7; ADKO, n=8). **k**, Insulin tolerant test (ITT) in mice from (**f**) on 19 weeks of HFD (Flox, n=7; ADKO, n=8). Data are mean ± s.e.m. *p<0.05, **p<0.01, ***p<0.001 and not significant (n.s.) by two-tailed Student’s t test (**b, d, e, i** and AUC in **j** and **k**) and two-way repeated measures ANOVA with post hoc test by Fisher’s LSD test (**c, f, j** and **k**).

### The ADKO mice are prone to HFD-induced obesity and abnormal glucose metabolism

To examine the physiological requirement of Breg for maintaining energy balance during energy overload, we challenged the ADKO mice with HFD. The ADKO mice consumed similar amounts food as control mice, yet gained significantly more body weight (Fig. 3f and Extended Data Fig. 5a), which was associated with substantially larger adipocytes with diminished UCP1 staining (Fig. 3g, h). EchoMRI measurement of a second cohort of ADKO mice confirmed that the body weight difference was due to fat mass, not lean mass (Fig. 3i and Extend Data Fig. 5b). We noted that the wild-type littermates of ADKO mice gained less body weight than wild-type littermates of *Breg* transgenic mice when fed HFD for a similar time period (Fig. 2g and Fig. 3f); this is likely due to their differences in genetic background and food intake. The ADKO mice and littermate controls consumed 2.6 grams per day (Extended data Fig. 5a), while the Breg transgenic mice and littermate controls consumed about 3.2 grams per day (Extended Data Fig. 3h). Remarkably, both steady-state and fasting glucose levels were higher, and glucose tolerance and insulin sensitivity were deteriorated in the ADKO mice fed on HFD for 18 and 19 weeks (Fig. 3j, k). Glucose tolerance and insulin tolerance tests were also performed at Week 5 and Week 6 of HFD feeding, respectively, when there was no difference in body weight compared with littermate controls, and similar results were obtained (Extend Data Fig. 5c, d). Thus, impaired beige adipocyte development caused by Breg ablation produced serious detrimental metabolic phenotypes that were opposite to those of Breg transgenic mice.

### Pharmacological levels of Breg can induce inguinal WAT browning through endocrine action

BAT-secreted Breg is unlikely to have significant contribution to WAT browning due to the substantially smaller mass of this depot. Indeed, neither BAT-selective overexpression nor BAT-selective deletion of *Breg* had any effect on inguinal WAT browning (Extended Data Fig. 6a, b). Therefore, the observed phenotypes of inguinal WAT thermogenesis in *Breg* transgenic mice and ADKO mice were likely due to paracrine signaling rather than endocrine signaling.

We next determined whether Breg is capable of functioning in an endocrine manner. We tail-vein infused adenoviruses expressing either *GFP*, mouse *Breg*, or human *BREG* into C57BL/6J wild type mice. As shown with human *BREG* adenovirus, this injection resulted in acute expression of BREG in the liver and its secretion into circulation with a concentration of about 0.45 µg/mL (Fig. 4a and Extended Data Fig. 7a). Importantly, no leaky expression of *BREG* was observed in the adipose tissue (Extended Data Fig. 7b). Circulating Breg led to a robust browning of inguinal WAT, as evidenced by strong induction of *Ucp1, Cidea*, mitochondrial genes and glycolytic genes, and occurrence of numerous smaller, Ucp1-positive adipocytes (Fig. 4b, c and Extended Data Fig. 7c), whereas no effect was observed in BAT and epididymal WAT (Extended Data Fig. 7d, e). In addition, expression of key metabolic genes in the liver was not altered (Extended Data Fig. 7f). Importantly, inguinal WAT browning was associated with lower steady-state glucose level, higher glucose tolerance, and increased glucose uptake (Fig. 4d, e). To examine potential benefits of Breg for glycemic control in an obese setting, we adenovirally expressed *Breg* in the liver of HFD-induced obese mice. Similar to what observed in the lean mice, obese mice receiving *Breg* adenovirus displayed a strong improvement of basal glucose level and glucose tolerance compared with obese mice receiving *GFP* adenovirus (Fig. 4f, g). Together, our data demonstrated that secreted Breg from liver acts in an endocrine manner to promote inguinal WAT browning and markedly improve glucose homeostasis.

**Fig. 4.**
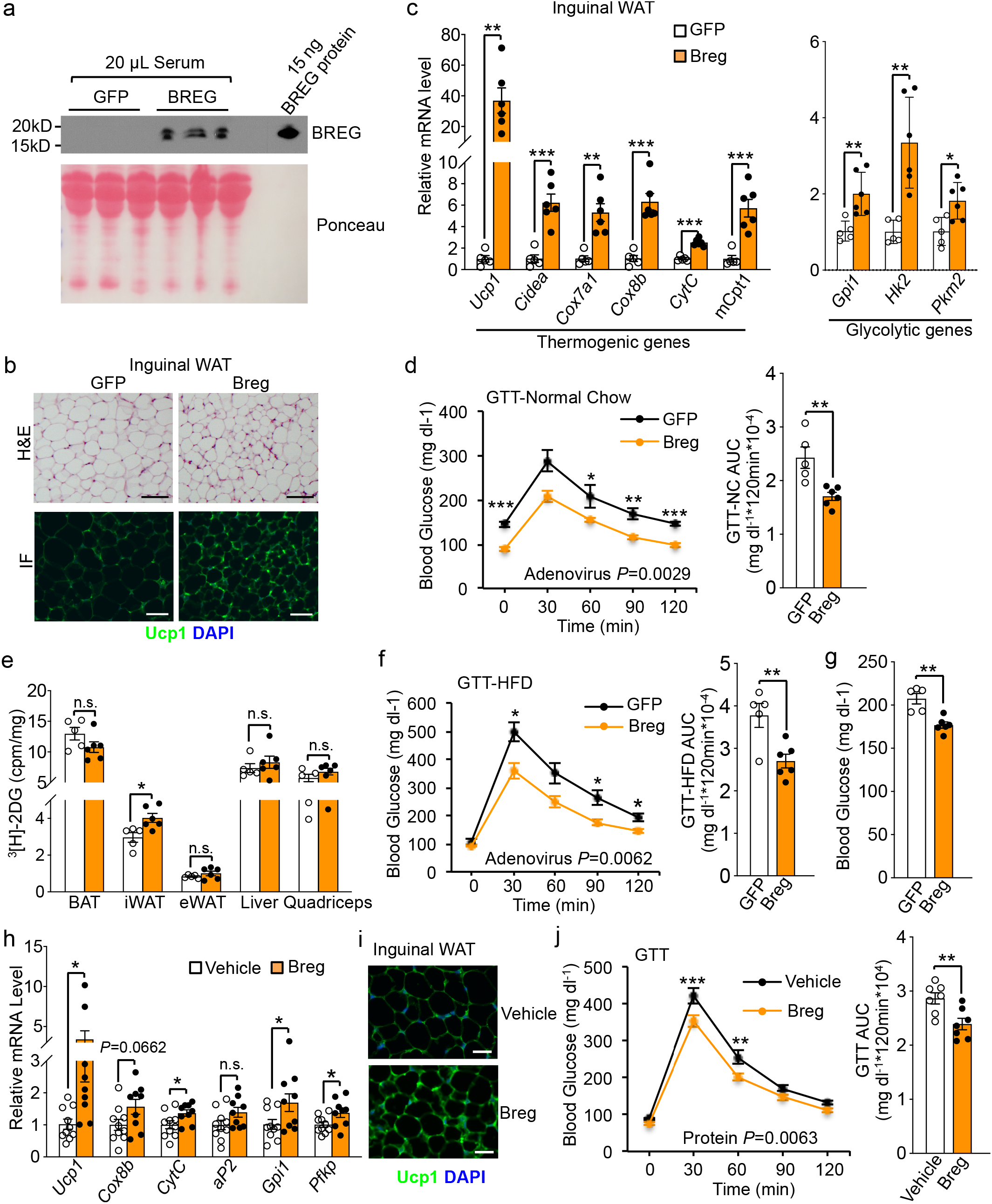
Endocrine action of Breg. **a**, Detection of circulating BREG protein. **b**, Representative images of H&E staining and Ucp1 immunofluorescence staining in iWAT (n=3 mice per group). Scale bar, 200 µm. **c**, mRNA expression of indicated genes in iWAT from adenovirus injected mice (GFP, n=5; Breg, n=6). **d**, GTT in mice fasted for 5 hours (GFP, n=5; Breg, n=6). **e**, Quantification of ^3^H-2DG uptake in indicated tissues from adenovirus injected mice (GFP, n=5; Breg, n=6) after 5 hours fasting. **f**, GTT in HFD-induced obese mice that were fasted overnight (GFP, n=5; Breg, n=6). **g**, Basal glucose of mice in (**f**) that were fasted for 5 hours (GFP, n=5; Breg, n=6). **h**, Gene expression in iWAT from WT mice intravenously injected with Breg protein or vehicle once a day for 9 days (Vehicle, n=10; Breg, n=9). **i**, Representative images of Ucp1 immunofluorescence staining in iWAT from mice in (**h)** (n=7 mice per group). Scale bar, 200 µm. **j**, GTT in another cohort of mice at day 7 of injection (n=7 mice per group). Data are mean ± s.e.m. *p<0.05, **p<0.01, ***p<0.001 and not significant (n.s.) by two-tailed Student’s t test (**c, e, g, h** and AUC in **d, f** and **j**) and two-way repeated measures ANOVA with post hoc test by Fisher’s LSD test (**d, f** and **j**).

We purified recombinant His-tagged Breg protein from HEK293 cells (Extended Data Fig. 7g). We then intravenously injected wild type mice with Breg protein daily for 9 days, which resulted in circulating level at 3.3 µg/mL. Compared with vehicle, Breg protein induced thermogenic gene expression and inguinal WAT browning (Fig. 4h, i and Extended Data Fig. 7h). Breg protein injected mice also exhibited improvement of glucose tolerance (Fig. 4j). We noted that recombinant Breg protein appeared to be less effective than adenoviral infusion; this may be due to the lengthy purification process that might affect Breg activity (Extended Data Fig. 7i). Our results provide additional evidence that pharmacological levels of Breg can induce inguinal WAT browning through endocrine signaling.

### Secreted Breg activates protein kinase A (PKA) signaling to exert its thermogenic function

We sought to determine the signaling pathway responsible for the thermogenic function of Breg. β3-AR and downstream PKA activation play a pivotal role in adipose thermogenesis^1, 2^. Our in vitro adipocyte culture data showing that Breg can induce *Ucp1* expression at basal condition and has an additive effect in the presence of β3-AR agonist (Extended Data Fig. 2c) indicate that β3-AR is unlikely to be involved. Indeed, *Ucp1* expression induced by adenoviral expression of Breg in adipocytes was not blocked by propranolol hydrochloride (β-blocker), a pan-β-receptor antagonist (Fig. 5a). To confirm this in vivo, *Breg* transgenic and control mice were administered with β-blocker. As expected, β-blocker markedly abolished *Ucp1* expression in inguinal WAT of control mice, but had no effect on *Ucp1* expression in *Breg* transgenic mice (Fig. 5b). Next, *Breg* transgenic mice and ADKO mice were housed at 30 °C and fed a normal chow diet for 1 month and 2 months, respectively. Differences in adipocyte morphology and Ucp1 level were observed in both inguinal WAT and BAT, compared with their respective littermate controls (Extended Data Fig. 8a, b, c, d). Although similar body weights as those of their respective littermate controls, WAT mass was lower in the *Breg* transgenic mice and was higher in the ADKO mice (Extended Data Fig. 8e, f). Thus, Breg-regulated thermogenesis and energy expenditure operates at thermoneutral condition. These results together suggest that Breg acts independently of β-AR. Interestingly, we found that induction of *Ucp1* expression by Breg was reduced by a merely 2-hr treatment with the PKA inhibitor H89 and was blocked by Melittin, an inhibitor for Gαs subunit of the heterotrimeric G protein (Fig. 5c, d), raising the idea that a G protein and PKA signaling axis might be responsible for the thermogenic activity of Breg. We thus examined PKA activation by western blot analysis using an antibody that detects a bulk of phosphorylated PKA substrates and an antibody against PKA-phosphorylated hormone sensitive lipase (pHSL). We found that phosphorylation of PKA substrates including HSL was increased in adipocyte culture infected with *Breg* adenovirus (Fig. 5e) as well as in inguinal WAT of *Breg* transgenic mice (Extended Data Fig. 9a), whereas inguinal WAT of ADKO mice had a decreased phosphorylation of PKA substrates and HSL (Fig. 5f). To provide independent evidence for PKA activation, we measured intracellular cAMP content. cAMP level was increased in primary inguinal adipocytes infected with *Breg* adenovirus (Extended Data Fig. 9b) and in inguinal WAT depot of *Breg* transgenic mice (Fig. 5g), and was decreased in inguinal WAT of ADKO mice (Fig. 5h). We next used recombinant Breg protein to directly examine PKA activation. Phosphorylation of PKA substrates in adipocyte culture was increased by purified Breg protein in a dose-dependent (Extended Data Fig. 9c) and time-dependent manner, with peak activation at 45-60 min (Fig. 5i). Moreover, phosphorylation of PKA substrates was induced in inguinal WAT of Breg protein administered mice (Extended Data Fig. 9d). These results together demonstrate that secreted Breg activates a G protein-cAMP-PKA signaling pathway.

**Fig. 5.**
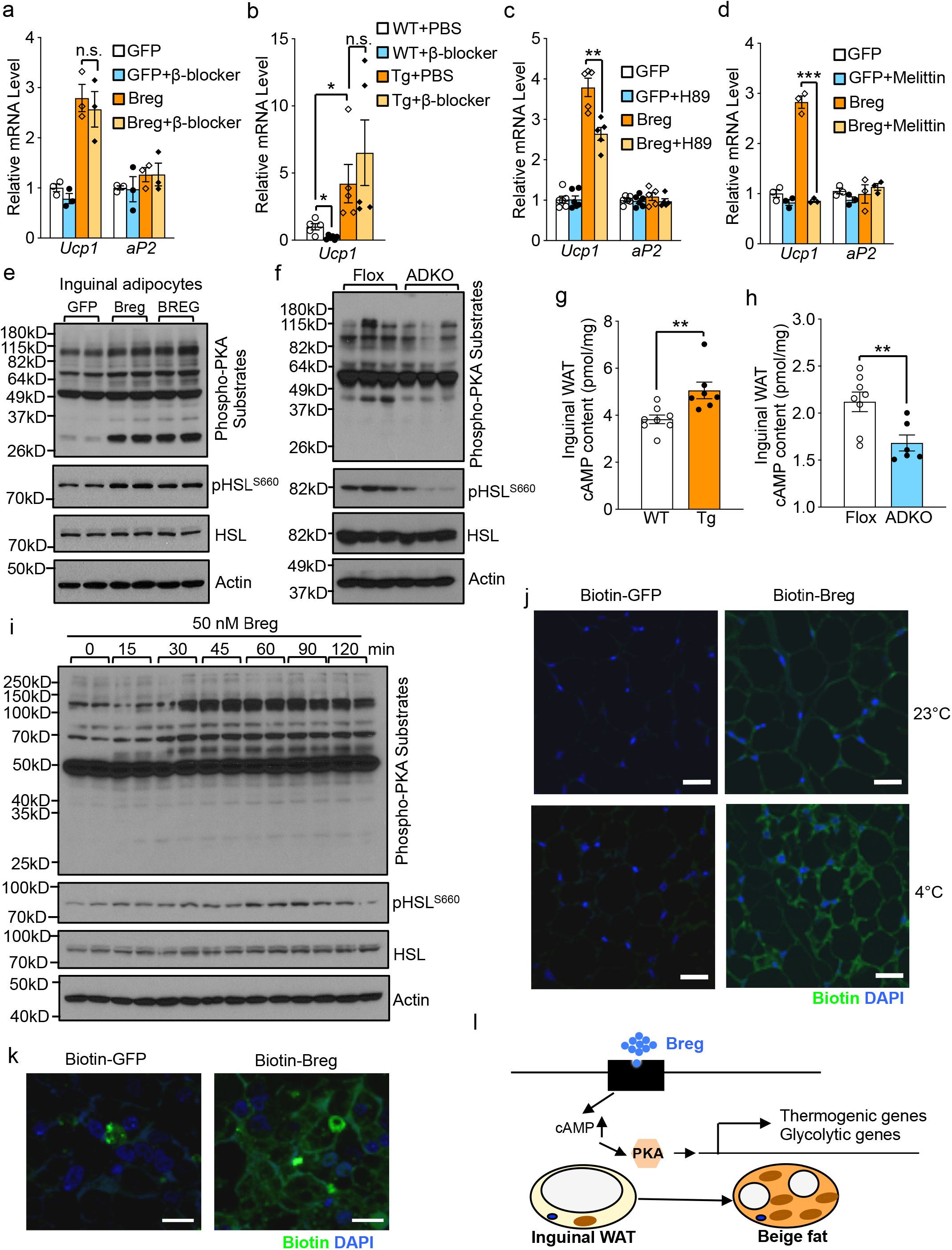
Breg functions through PKA signaling. **a**, Gene expression in inguinal adipocytes that overexpressing *Breg* and treated with β-blocker (100 nM) for 24 hours (n=3). **b**, *Ucp1* expression in iWAT from male *Breg* Tg mice and WT controls after administrated with β-blocker (n=5-7). **c, d**, Gene expression in inguinal adipocytes overexpressing *Breg* and treated with 30 µM H89 (n=5-6) for 2 hours (**c**) or 1 µM Melittin (n=3) for 24 hours (**d**). **e**, Western blot of phosphorylated PKA substrates and HSL in inguinal adipocytes transduced with indicated adenovirus. **f**, Western blot of phosphorylated PKA substrates and HSL in iWAT from female *Breg* ADKO mice and Flox controls after 3 hours cold exposure. **g**, cAMP levels in iWAT from male *Breg* Tg mice (n=8) and WT controls (n=7). **h**, cAMP levels in iWAT from mice in (**f**) (Flox, n=8; ADKO, n=6). **i**, Western blot of phosphorylated PKA substrates and HSL in brown adipocytes treated with Breg protein. **j**, Binding of biotin labeled-Breg to iWAT from WT mice housed at 23°C or cold (4°C) challenged for 8 hours (n=3 mice per group). Scale bar, 20 µm. **k**, Binding of biotin labeled-Breg to brown adipocyte surface. Scale bar, 20 µm. **l**, Working model of Breg. Data are mean ± s.e.m. *p<0.05, **p<0.01, ***p<0.001 and not significant (n.s.) by two-tailed Student’s t test.

Our data indicate that Breg likely binds to a cell surface receptor distinct from β-AR to activate downstream signaling in adipocytes. To provide evidence for the existence of such a receptor, conditioned medium from HEK293 cells expressing Breg fused to the C-terminus of secreted alkaline phosphatase (SEAP) was incubated with frozen tissue slices. Binding of SEAP-Breg was determined by examining alkaline phosphatase activity. Whereas liver, skeletal muscle and brain had a near background binding signal, a specific binding of Breg to adipose tissue sections was detected, which can be competed away by incubation with Breg-containing conditioned medium (Extended Data Fig. 9e, f). In a second set of experiments, recombinant Breg protein was labeled with biotin, incubated with inguinal WAT tissue sections, and stained with Alexa Fluor 488-conjugated Streptavidin. We found that Breg, not GFP control, bound to adipose sections (Fig. 5j). Interestingly, the binding was increased in adipose tissue of mice challenged by cold (Fig. 5j). Breg was also bound to cell surface of cultured adipocytes (Fig. 5k). Together, our data suggest that Breg activates PKA signaling through binding to a putative receptor at adipocyte surface (Fig. 5l).

## Discussion

Our study shows that Breg, encoded by a previously uncharacterized gene, is a bona fide, BAT-enriched adipokine. While a few batokines were previously identified, they are highly expressed in other tissues and/or have a very low expression in adipose tissue, and the importance of endogenous levels of these proteins secreted from adipose tissue was unknown^12^. Breg is almost exclusively expressed in adipose tissue and, based on RNA-Seq data, *Breg* mRNA is one of the most abundant transcripts in BAT. Adipose-specific *Breg* deletion decreases cAMP content and PKA activity, suppresses inguinal WAT browning, and leads to HFD induced obesity and hyperglycemia. Thus, Breg is one of the few BAT-enriched adipokines known to date and plays an indispensible role in metabolic physiology^29^. Currently we were not able to measure endogenous, circulating Breg level, which requires a more sensitive assay. In this regard, it is important to note that our data suggest that endogenous Breg regulates inguinal WAT thermogenesis through paracrine signaling, and BAT-secreted Breg has little contribution given the substantially smaller mass of this fat depot. Yet, when administered at a pharmacological level in circulation, Breg can have a robust effect through endocrine action.

The mechanisms underlying the beneficial effect of WAT browning on glucose metabolism were unclear^8^. One important finding in our study is that Breg markedly improves glucose homeostasis independent of body weight. Recent work by others has shown that mild cold promotes glycolytic beige fat formation when β-adrenergic signaling is blocked^30^. We found that inguinal WAT browning promoted by Breg is also associated with increased expression of glycolytic genes, and interestingly, this increase can occur at room temperature and without blocking β-adrenergic signaling. The glucose-lowering effect of Breg could be in part attributed to glycolysis. Alternatively, Breg may additionally have a signaling role in glycemic control. It has been postulated that beige fat secretes factors to directly regulate glucose metabolism^31, 32^.

Breg binds to a putative cell surface receptor and subsequently activates downstream PKA to promote WAT browning and metabolic healthy independently of β-AR activation. While the PKA signaling is also required in BAT, the effect of Breg was only manifested under a thermoneutral condition, not at ambient temperature. At ambient temperature, BAT has a constitutively active, basal β3-AR signaling and hence high basal PKA activity due to its abundant sympathetic innervation, which likely renders Breg less important. On the other hand, in the inguinal WAT, activation of both pathways, converging on downstream PKA activation, appears to have a synergistic effect, suggesting a concerted action. This is also consistent with our observations that acute β3-AR activation stimulates Breg secretion and chronic β3-AR activation increases Breg expression. Thus, Breg serves to further ensure a full thermogenic response in inguinal WAT. In summary, our study provides novel molecular insights into our understating of signaling pathways underlying adipose thermogenesis and suggests a therapeutic potential of Breg for the treatment of obesity and diabetes.

## Methods

### Adipocyte culture

Immortalized brown preadipocytes were generated previously^33^. At day -2 of differentiation, 70% confluent brown preadipocytes were cultured in DMEM medium (Gibco) containing 10% FBS (Atlanta Biologicals) and supplemented with 20 nM insulin (Sigma) and 1 nM triiodotyronin (Sigma) (differentiation medium). At day 0, differentiation was induced by culturing cells in differentiation medium supplemented with 0.5 mM isobutylmethylxanthine (Acros organics), 0.5 µM dexamethasone (Acros organics), and 0.125 mM indomethacin (Alfa Aesar) for 2 days. Cells were then changed back to differentiation medium and fresh medium was replenished every 2 days. Inguinal stromal-vascular fractions (SVF) were isolated from 2-week-old C57BL/6J mice with collagenase D (Roche) digestion and differentiated as described previously^34^. In brief, inguinal preadipocytes differentiation was started by culturing confluent cells in DMEM/F12 (Gibco) medium containing 10% FBS, 850 nM insulin, 1 nM triiodotyronin, 0.5 mM isobutylmethylxanthine, 0.5 µM dexamethasone, and 0.125 mM indomethacin. After 2 days, cells were cultured in DMEM/F12 medium containing 10% FBS, 850 nM insulin and 1 nM triiodotyronin and cell medium was changed every 2 days. For conditioned medium experiment, after removal of small molecules with a 10-kDa cut-off filter, serum-free conditioned medium collected from immortalized brown adipocyte culture was used to treat primary inguinal adipocytes. Heat-treated medium was heated at 95°C for 10 minutes. Both brown and inguinal adipocyte were fully differentiated and harvested at day 6. In some experiments, differentiated adipocytes were treated with 10 µM CL-316,243 (R&D Systems) as indicated.

### Identification of brown fat-enriched adipokines

Proteomics of proteins secreted from immortalized brown adipocytes was done by MS Bioworks (Ann Arbor, MI). 350 µg total protein concentrated with 3-kDa cut-off centrifugal filter (Millipore) from serum-free medium was subjected to SDS-PAGE with multiple band excision and in-gel digestion. Liquid Chromatography with tandem mass spectrometry (LC-MS/MS) was performed. Protein identification was performed with Mascot. Protein visualization and validation was performed with Scaffold. We screened previously published RNA-seq dataset (GSE56367)^13^ and generated a list of genes abundantly expressed in BAT (FPKM>40) that are >3-fold and >5-fold relative to epididymal WAT and skeletal muscle, respectively. Overlapping genes from the above analyses were then predicted for secretion potential using SignalP and SecretomeP (www.cbs.dtu.dk/services)^17,35^ and the Universal Protein Resource (UniProt).

### Generation of *Breg*-*flag* knock-in cells

CRISPR/Cas9 system was used to knock-in a Flag tag immediately before the stop codon at the endogenous *Breg* locus in immortalized brown preadipocytes. Oligonucleotides (F: CACCGCTCGGCTTCGACTGAGGCC, R: AAACGGCCTCAGTCGAAGCCGAGC) were cloned into pX330-puro vector^36^ for expression of the guide RNA. A 1-kb flanking *Breg* genomic fragment with the Flag tag inserted before the stop codon was produced by PCR and cloned into pBluescript II vector. The above plasmids were co-transfected into brown preadipocytes and selected with puromycin (Sigma). Cells were re-plated without puromycin and single cell clones were examined by genotyping. Correctly targeted single cell clones were further verified by sequencing of produced *Breg* transcript.

### Animals

C57BL/6J wild type mice (Stock No. 000664) were obtained from the Jackson Laboratory. 3-month-old male C57BL/6J mice were used for adenoviruses or protein injection. *Ucp1* knockout mice were obtained from David A. Guertin lab. *Breg* transgenic mice were generated by core facility of UMASS Medical School. *Breg* cDNA was fused downstream of the 5.4 kb *aP2* promoter. The transgenic DNA fragment was gel-purified and injected into fertilized embryos harvested from C57BL/6J×SJL hybrid mice. Transgenic lines were backcrossed with C57BL/6J for at least three generations. To generate *Breg* knockout mice, *Breg* embryonic stem cell (ESC) clones (in C57BL/6N background) with conditional potential were obtained from European Mouse Mutant Cell Repository. Germ-line transmissible mice were generated by core facility of UMASS Medical School. These mice were then crossed with *Flp* mice (Stock No. 012930, Jackson Laboratory) to remove the targeting cassette, resulting in exon 3 flanked by loxP sites. The Floxed mice were crossed with *adipnectin*-*cre* mice^28^ to generate adipose-specific *Breg* knockout (ADKO) mice.

Mice were maintained under a 12 hours light/12 hours dark cycle at constant temperature (23°C) with free access to food and water. Normal diet containing 4% (w/w) fat and high-fat diet containing 35% (w/w) fat were purchased from Harlan Teklad and Bioserv, respectively. Daily food intake of single caged, 2- to 3-month-old mice was measured for 2 weeks. Fat mass and lean mass were measured by Body Composition Analyzer EchoMRI (Echo Medical Systems). β3-AR agonist CL-316,243 (R&D Systems) was intraperitoneally injected into mice at 1mg/kg body weight for 2 days. β-blocker (Sigma) was intraperitoneally injected into mice at 25 mg/kg body weight once a day for 7 days. For acute cold exposure, mice were placed at 4°C with water but without food, and core body temperature was measured with the Microtherma 2 rectal probe (Thermoworks). For studies on thermoneutral condition, mice were maintained under a 12 hours light/12 hours dark cycle at 30°C with free access to food and water. Gender-matched littermate controls were used in all the studies. All animal studies were performed according to procedures approved by the UMASS Medical School’s Institutional Animal Care and Use Committee (IACUC).

### Adenovirus production, purification, and injection

Full-length mouse *Breg* and human *BREG* cDNA were generated by PCR. Adenoviral *Breg* overexpression plasmids were constructed using AdEasy-1 system^37^. Adenoviruses were purified with cesium chloride ultracentrifugation. All viral titers were predetermined, and same number of viral Plaque Formation Unit (PFU) was used for experimental and control samples.

For tail-vein injections, *GFP*, mouse *Breg* or human *BREG* adenoviruses (1-2×10^10^ PFU/mice) were injected in a total volume of 150 µL. To overexpress *Breg* in BAT, *Breg* or *GFP* adenoviruses (1×10^10^ PFU at the volume of 20 µL per site) were injected into six different sites of bilateral BAT of 3-month-old C57BL/6J male mice to cover the whole fat pad. To delete *Breg* in BAT, *Cre* or *GFP* adenoviruses (1×10^10^ PFU at the volume of 20 µL per site) were injected into six different sites of bilateral BAT of 6-month-old *Breg* floxed female mice to cover the whole fat pad. To overexpress *Breg* in iWAT, *Breg* and *GFP* adenoviruses (1×10^10^ PFU in 30 µl per site) were injected into the left and right iWAT pad of the same C57BL/6J male mice, respectively. Five different sites were injected to cover the whole fat pad. One week after the injection, the mice were dissected and tissues were harvested.

### Gene expression analysis

Total RNA was extracted from cells or tissues using TRIzol reagent (Invitrogen). 1 µg total RNA was converted into first strand complementary DNA (cDNA) with random primers using an IScript cDNA synthesis kit (Invitrogen). qPCR was performed with SYBR green fluorescent Dye (Bio-Rad) using an ABI7300 PCR machine, and *36B4* was used as an internal control. Relative mRNA expression was determined by the ΔΔ-Ct method. The sequences of qPCR primers used in this study are available upon request.

### Protein extraction and western blotting

Total protein was extracted from cells or adipose tissues by using lysis buffer [100 mM NaCl, 50 mM Tris (pH7.5), 0.5% Triton X-100, 5% (w/v) glycerol] supplemented with complete protease inhibitor cocktail (Roche), phosphatase inhibitor cocktail (Sigma) and phenylmethylsulfonyl fluoride (PMSF) (Thermo Fisher Scientific). For secretion assay of Breg, *Breg* adenovirus-transduced brown adipocytes were cultured in serum-free medium with 1× protein transport inhibitor cocktail (10.6 µM Brefeldin A and 2 µM Monensin) (Fisher scientific, #50-930-9), 10 µM oligomycin, or 10 µM FCCP (Carbonyl cyanide-4 (trifluoromethoxy) phenylhydrazone) for indicated times, and conditioned medium was collected. For secretion assay of Breg with CL316,243, Breg adenovirus-transduced brown adipocytes were cultured in serum-free medium with 10 µM CL316,243. Medium was harvested at indicated time intervals and fresh serum-free medium with 10 µM CL316,243 was supplied. The medium was concentrated with a 3-kDa cut-off centrifugal filter (Millipore). Samples were subjected to SDS-PAGE under reducing conditions, transferred, and immunoblotted with antibodies against Flag (Sigma, #F7425), Breg (MyBioSourse, #MBS1493234), Ucp1 (Sigma, #U6382), Phospho-PKA Substrate (Cell signaling, #9624), HSL (Cell Signaling, #4107), pHSL^S660^ (Cell Signaling, #4126), Tubulin (DSHB, #E7), Adiponectin (Cell Signaling, #2789), or Actin (Santa Cruz, #sc-47778).

For serum samples, Albumin/IgG was removed from 60 µL serum according to the manufacturer’s instructions (Millipore). The albumin/IgG-depleted serum was concentrated to 60 µL with 10-kDa cut-off centrifugal filter. 30 µL serum was subjected to SDS-PAGE under reducing conditions, transferred, and immunoblotted with antibodies against Breg (MyBioSourse, #MBS1493234), ponceau S staining as loading control.

### Glucose and insulin tolerance tests

For glucose tolerance test, mice were fasted overnight or otherwise indicated. Glucose (2 g/kg body weight) was administered intraperitoneally, and blood glucose was measured at 0, 30, 60, 90, and 120 min. For insulin tolerance test, mice were fasted 5 hours. Insulin (0.75 U/kg body weight) was administered intraperitoneally, and blood glucose was measured at 0, 30, 60, 90, and 120 min.

### Oxygen consumption assay

Subcutaneous inguinal WAT depots were weighed and chopped into small pieces. Portions with equal amount (50 mg) were suspended in 5 mL Dulbecco’s phosphate-buffered saline supplemented with 25 mM glucose, 1 mM pyruvate and 2% BSA, and respiration was measured with a biological oxygen monitor (YSI model 5300A).

### *In vivo* Glucose uptake

In vivo glucose uptake was measured as previously described^38^. Briefly, wild-type mice were tail-vein infused with *GFP* or *Breg* adenoviruses, and at day 7, mice were fasted for 5 hr. *Breg* floxed and ADKO mice were cold challenged for 3 hr without food. ^3^H-2-deoxyglucose (100 μCi/kg body weight) (PerkinElmer) was then intravenous injected. One hour post injection, mice were sacrificed and tissues were collected, weighed and snap frozen in liquid nitrogen. Tissues were digested by incubating in 500 µL of 1 M NaOH at 60°C for 60 min and neutralized with 500 µL of 1 M HCl. 200 µL of this neutralized solution were added to 1 mL of 6% perchloric acid, vortexed and centrifuged at 13,000 g for 5 min. 800 µL supernatant was mixed with 5 mL scintillation cocktail and total radioactivity was quantified in c.p.m. by liquid scintillation counting.

### Protein purification and injection

To increase the yield and facilitate the purification of secreted Breg, we added a signal peptide at the N-terminus of Breg and a 6×His tag at its C-terminus using pHL-sec vector (Addgene)^39^. The resulting DNA fragment was cloned into pENTR1A vector (Addgene). Lentiviral construct was generated via recombination of pENTR1A-Breg into pLenti CMV/TO Puro DEST (Addgene), and lentiviruses were produced by co-transfection along with plasmids pLP1, pLP2, and pVSVG into HEK293 cells. HEK293 cells were infected with lentiviruses and Breg expressing stable cells were selected with puromycin. Serum-free medium from stable cells were collected, and incubated with Ni-NTA Agarose column at 4°C for 2 hours. After washing with buffer (PBS containing 5 mM imidazole), Breg protein was eluted with buffer (PBS containing 250 mM imidazole), concentrated and then buffered in PBS. Endotoxin level (0.08 EU/ml) was measured using a commercial kit (Thermo Scientific, #88282). Purified Breg protein was used to treat adipocytes, or intravenously inject into 3-month-old C57BL/6J male mice (1 mg/kg body weight) in a total volume of 150 µL. PBS was used as vehicle control. Breg protein was used within 1 week after purification.

### Competitive ELISA assay

The competitive ELISA assay was modified from previously described^40^. Wells of microplate (R&D, #DY990) were filled with 50 µL of recombinant Breg protein (250 ng/mL) in coating buffer (PBS, pH=7.4), with the exception of 2 blank controls filled with coating buffer, and were incubated overnight at 4°C. They were then washed twice with washing buffer (PBS with 0.05% Tween-20), and filled with 200 µL blocking buffer (PBS with 3% BSA) and incubated for 2 hr at room temperature. 130 µL of purified mouse Breg protein with various concentrations or samples were preincubated with Breg antibody (250 ng/mL) for l hr at room temperature. Wells were washed twice with washing buffer and filled with 50 µL per well of preincubated standards or samples in duplicate and incubated for 3 hr at room temperature. After washed three times with washing buffer, 50 µL of the HRP-conjugated mouse-anti-rabbit antibody (1:5000) was added to all the wells and incubated for 1 hr at room temperature. Then they were washed three times with washing buffer, filled with 100 µL TMB substrate solution (Thermo Fisher Scientific, #34028) and incubated for 30 min in the dark at room temperature. The reaction was terminated by the addition of 100 µL stop solution (2 M sulfuric acid) per well. Optical densities at 450 nm were measured on an ELISA plate reader within 10 min. All dilutions were done in the blocking buffer.

### cAMP assay

cAMP assay was done according to the manufacturer’s instructions (R&D Systems, #KGE012). In brief, inguinal WAT depot was weighed, and equivalent amount (~50 mg) was minced with scissors and then homogenized in 500 µL 0.1 N HCl. Lysates were collected and centrifuged at 10,000 g for 10 min. Then 200 µL supernatant was transferred to new tubes and mixed with 28 µL 1 N NaOH and diluted with calibrator diluent RD5-55. Samples were centrifuged again at 10,000 g for 10 min, and 100 µL supernatant was used for assay. To measure cAMP in cell culture, cells were resuspended in cold 0.1 N HCl/Cell Lysis Buffer after washed with cold PBS, incubated at room temperature for 10 min and neutralized with 1 N NaOH (1:10). Cell lysates were centrifuged at 600 g for 10 min, diluted 2-fold with calibrator diluent RD5-55 and 100 µL supernatant was used for assay.

### Breg binding assay

Recombinant Breg was purified from conditioned medium of HEK293 cells, and recombinant GFP protein was purified from *E. coli* BL21. Recombinant Breg and GFP protein was labeled with membrane non-permeable biotin according to the manufacturer’s instructions (Thermo Fisher Scientific).

Three-month-old wild type male mice were challenged with or without cold (4 °C) for 8 hr, and then the mice were sacrificed. The inguinal WAT were fixed with 10% formalin overnight and paraffin-embedded. The formalin-fixed and paraffin-embedded tissue sections were deparaffinized and rehydrated by graded concentrations of ethanol solutions. The sections were incubated with biotin-GFP or biotin-Breg protein (500 ng/ml) at room temperature for 1 hr and gently washed by PBS. Next, the tissue sections were incubated with Alexa Fluor 488-conjugated streptavidin (Invitrogen, #S32354) (1:500 dilution) and DAPI (Sigma, #D9542) at room temperature for 2 hr and gently washed by PBS. The images were acquired with an inverted Nikon Eclipse Ti2 confocal microscope (Nikon Instruments/Nikon Corp).

In vitro differentiated brown adipocytes were treated with biotin-GFP or biotin-Breg protein (500 ng/ml) at 37 °C for 1 hr. The cells were gently washed by PBS and fixed with 3.7% formalin at 37 °C for 15 min without permeabilization. Subsequently, the cells were incubated with Alexa Fluor 488-conjugated streptavidin (Invitrogen, #S32354) (1:500 dilution) at room temperature for 2 hr. Then, the cells were stained with DAPI (Sigma, #D9542) at room temperature for 20 min and gently washed by PBS. The images were acquired with an inverted Nikon Eclipse Ti2 confocal microscope (Nikon Instruments/Nikon Corp).

Breg tissue binding assay through detecting SEAP activity was performed as previously described^29, 41^. Briefly, HEK293 cells were transfected with plasmids expressing SEAP or SEAP-Breg. Serum-free medium was collected and concentrated using 10-kDa centrifugal filter. Frozen tissue slices were incubated with SEAP or SEAP-Breg conditioned medium for 90 min at room temperature. They were fixed in a solution containing 20 mM HEPES (pH 7.4), 60% acetone and 3% formaldehyde after washed four times with PBS containing 0.1% Tween-20. After inactivating endogenous alkaline phosphatase at 65 °C for 30 min, the enzymatic activity derived from the fusion protein was detected using NBT/BCIP substrate (Sigma). For competition binding, frozen tissue slices were pre-incubated for 60 min with 5-fold concentrated conditioned medium from HEK293 cells expressing Breg.

### Immunofluorescence staining and histology

Differentiated brown adipocytes were gently washed with DMEM and treated with PBS or CL-316, 243 (10 µM) for 45 min. Cells were then fixed with 3.7% formalin at 37 °C for 15 min. The cells were permeabilized with ice-cold 100% methanol at -20 °C for 10 min and washed twice with PBS. Next, the cells were incubated with blocking buffer (PBS containing 5% normal goat serum and 0.3% Triton X-100) at room temperature for 60 min and incubated with Breg antibody (1:200 dilution) and β-actin antibody (1:200 dilution) in blocking buffer at 4 °C overnight. After incubation, the cells were gently washed twice with PBS and incubated with Alexa Fluor 488-conjugated (Thermo Fisher, #A-11034) and 594-conjugated secondary antibody (Thermo Fisher, #A-11032) at room temperature for 2 hr. Finally, the cells were stained with DAPI (Sigma, #D9542) at room temperature for 20 min, and images were acquired with an inverted Nikon Eclipse Ti2 confocal microscope (Nikon Instruments/Nikon Corp).

Tissues fixed with 10% formalin were paraffin-embedded. H&E staining was performed according to standard procedures. Immunofluorescence staining was done as described previously^42^. Paraffin-embedded tissue sections were deparaffinized and rehydrated through graded ethanol solutions. Slides were incubated with Ucp1 antibody (Sigma, 1:500) in dilution buffer (PBS containing 1% BSA and 0.3% Triton X-100) at 4°C overnight after pre-incubated with blocking buffer (PBS containing 5% normal goat serum and 0.3% Triton X-100) for 1 hr. Next, the slides were washed 3 times with PBS and incubated with Alexa Flour 488-conjugated goat anti-rabbit secondary antibody (Thermo Fisher Scientific) for 2 hr at room temperature. The slides were then stained with DAPI (Sigma) for 1 hr after washed 3 times with PBS. Images were acquired and processed with the same settings.

### Statistical analysis

Data are presented as mean ± s.e.m. Two-tailed Student’s t test and two-way repeated measures ANOVA with post hoc test were used for statistical analysis. p < 0.05 was considered statistically significant in all the experiments. The statistical parameters and the number of mice used per experiment are stated in the Figure legends.

### Data availability

The data that support the findings of this study are available from the corresponding author upon reasonable request.

## Acknowledgments

We thank the Transgenic Animal Modeling Core at University of Massachusetts Medical School for the generation of *Breg* transgenic and knockout mice. We thank the Morphological Core at University of Massachusetts Medical School for technical help. We thank Dr. Tom Fazzio for pX330-puro plasmid, Dr. Tingting Huang for construction of *Breg*-*Flag* knock-in plasmids, Sasha Lee for help with the use of EchoMRI, Danning Li for help on writing of this manuscript. This work was supported by NIH/NIDDK (R01DK115918 and R01DK116872) and the Early Exploration Award from Novo Nordisk.

## Author Contributions

Q.C., L.H., and D.P. designed and performed the experiments, and analyzed the data. Y.-X.W. designed the experiments and analyzed the data. K.H., R.L., and L.J.Z. performed bioinformatics analysis. D.A.G. provided the *Ucp1* knockout mice. Q.C. and Y.-X.W. wrote the manuscript.

## Additional information

Supplementary Information is available for this paper. Correspondence and requests for materials should be addressed to Y.-X.W.

