## Supplemental figures for "A novel batokine Breg controls adipose thermogenesis and glucose homeostasis"

### Extended Data Table 1

**BAT-enriched proteins that are also present in conditioned medium of brown adipocyte culture.**

| Gene | Description |
|---|---|
| Aldoa | aldolase A, fructose-bisphosphate |
| Prkar2b | protein kinase, cAMP dependent regulatory, type II beta |
| Apoc1 | apolipoprotein C-I |
| Tkt | transketolase |
| Agpat2 | 1-acylglycerol-3-phosphate O-acyltransferase 2 (lysophosphatidic acid acyltransferase, beta) |
| Thrsp | thyroid hormone responsive |
| Acly | ATP citrate lyase |
| Mecr | mitochondrial trans-2-enoyl-CoA reductase |
| Taldo1 | transaldolase 1 |
| Acadl | acyl-Coenzyme A dehydrogenase, long-chain |
| Hadhb | hydroxyacyl-Coenzyme A dehydrogenase/3-ketoacyl-Coenzyme A thiolase/enoyl-Coenzyme A hydratase (trifunctional protein), beta subunit |
| Echs1 | enoyl Coenzyme A hydratase, short chain, 1, mitochondrial |
| Hadh | hydroxyacyl-Coenzyme A dehydrogenase |
| Hibch | 3-hydroxyisobutyryl-Coenzyme A hydrolase |
| Decr1 | 2,4-dienoyl CoA reductase 1, mitochondrial |
| Acsl1 | acyl-CoA synthetase long-chain family member 1 |
| Etfa | electron transferring flavoprotein, alpha polypeptide |
| Slc16a1 | solute carrier family 16 (monocarboxylic acid transporters), member 1 |
| Acadm | acyl-Coenzyme A dehydrogenase, medium chain |
| Dhrs4 | dehydrogenase/reductase (SDR family) member 4 |
| Nudt7 | nudix (nucleoside diphosphate linked moiety X)-type motif 7 |

(continued)

| **Gene** | **Description** |
|---|---|
| Ddt | D-dopachrome tautomerase |
| Atpaf2 | ATP synthase mitochondrial F1 complex assembly factor 2 |
| Eci1 | enoyl-Coenzyme A delta isomerase 1 |
| Gpd1 | glycerol-3-phosphate dehydrogenase 1 (soluble) |
| Bckdha | branched chain ketoacid dehydrogenase E1, alpha polypeptide |
| Aifm1 | apoptosis-inducing factor, mitochondrion-associated 1 |
| Zadh2 | zinc binding alcohol dehydrogenase, domain containing 2 |
| Acads | acyl-Coenzyme A dehydrogenase, short chain |
| Dlst | dihydrolipoamide S-succinyltransferase (E2 component of 2-oxo-glutarate complex) |
| Acaca | acetyl-Coenzyme A carboxylase alpha |
| **1700037H04Rik** | **RIKEN cDNA 1700037H04 gene** |
| Pfkl | phosphofructokinase, liver, B-type |
| Acat2 | acetyl-Coenzyme A acetyltransferase 2 |
| Uck1 | uridine-cytidine kinase 1 |
| Fam82a2 | Rmdn3 |
| Eno1 | enolase 1, alpha non-neuron |
| Ahcyl1 | S-adenosylhomocysteine hydrolase-like 1 |
| Coasy | Coenzyme A synthase |
| Gngt2 | guanine nucleotide binding protein (G protein), gamma transducing activity polypeptide 2 |
| Acaa2 | acetyl-Coenzyme A acyltransferase 2 (mitochondrial 3-oxoacyl-Coenzyme A thiolase) |
| 4931406C07Rik | RIKEN cDNA 4931406C07 gene |

### Extended Data Fig. 1

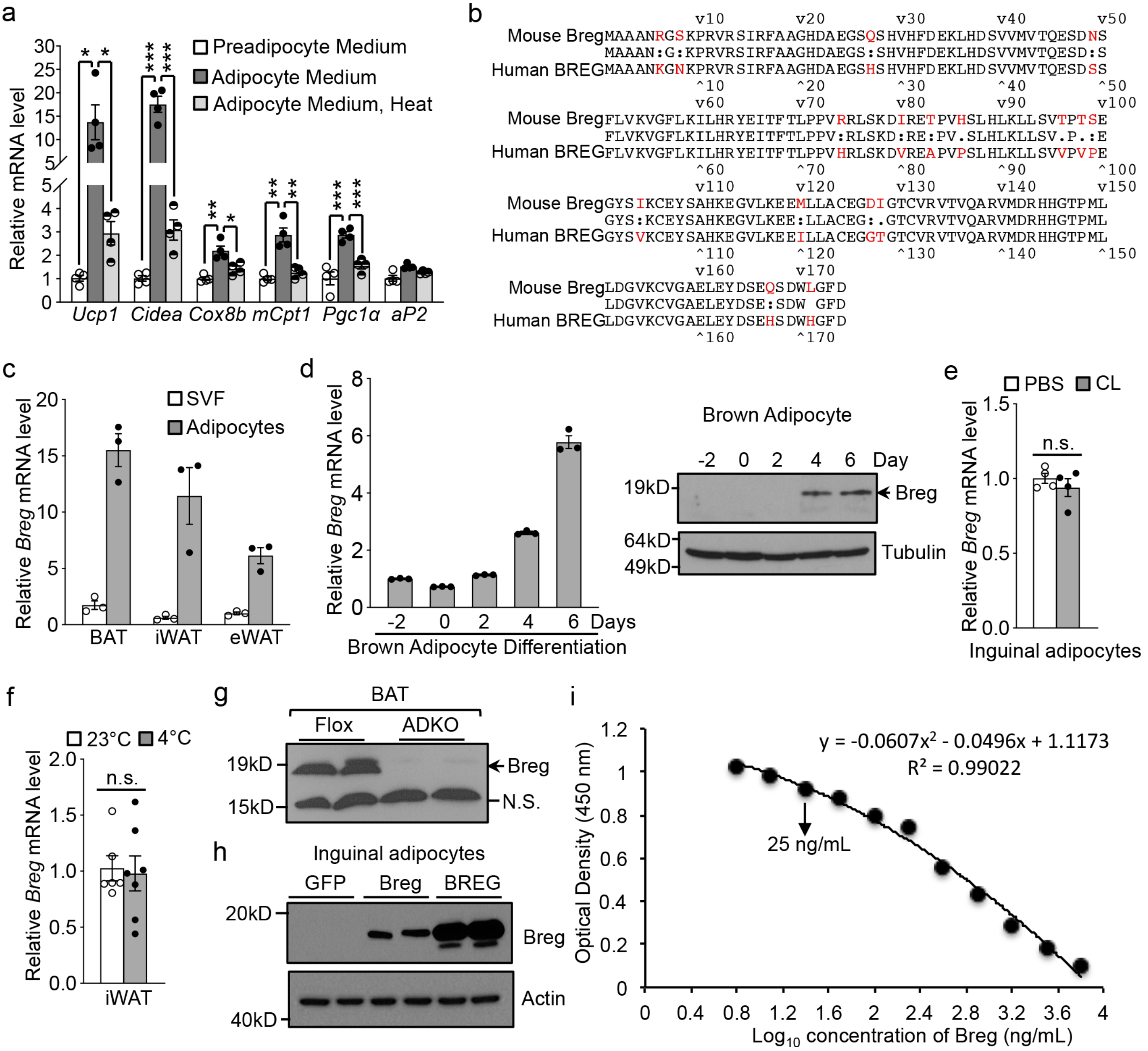


**Extended Data Fig. 1 | Identification of Breg. a**, Gene expression in inguinal adipocytes treated with indicated medium for 6 days (n=4). **b**, Alignment of mouse Breg and human BREG protein sequences. **c**, *Breg* mRNA expression in stromal vascular fraction (SVF) and adipocyte fraction (n=3 mice). **d**, *Breg* mRNA (n=3) and protein levels during brown adipogenesis. **e**, *Breg* expression in primary inguinal adipocytes treated with 10 μM CL-316,243 for 3 hours (n=4). **f**, *Breg* expression in iWAT from WT male mice housed at 23°C (n=5 mice) or 4°C (n=7 mice) for 6 hours. **g**, Western blot of Breg in BAT from *Breg* adipose tissue knockout mice (ADKO) and Flox controls. N.S., non-specific band. **h**, Western blot of Breg in inguinal adipocytes transduced with *GFP*, *Breg* or *BREG* adenoviruses. **i**, Standard curve of the competitive ELISA assay for Breg. Data are mean ± s.e.m. n.s., not significant by two-tailed Student’s t test.

Extended Data Fig. 2

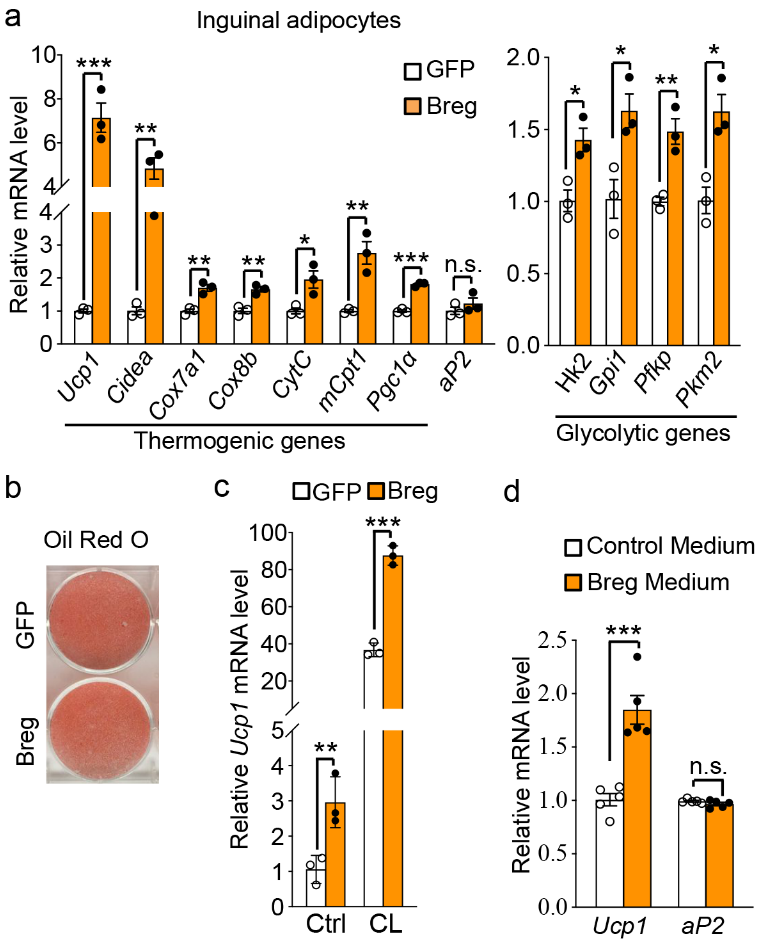


**Extended Data Fig. 2 | Breg induces thermogenic gene expression in adipocytes.** **a**, Gene expression analysis in primary inguinal adipocytes transduced with *Breg* or *GFP* adenoviruses (n=3). **b**, Oil Red O staining of inguinal adipocytes in (**a**). **c**, Gene expression analysis in inguinal adipocytes transduced with *Breg* or *GFP* adenoviruses (n=3) treated with PBS (Ctrl) or 10 μM CL-316,243 (CL) for 3 hours. **d**, Gene expression analysis in inguinal adipocytes treated with indicated conditioned medium for 6 days (n=5). Data are mean ± s.e.m. *p<0.05, **p<0.01, ***p<0.001 and not significant (n.s.) by two-tailed Student's t test.

Extended Data Fig. 3

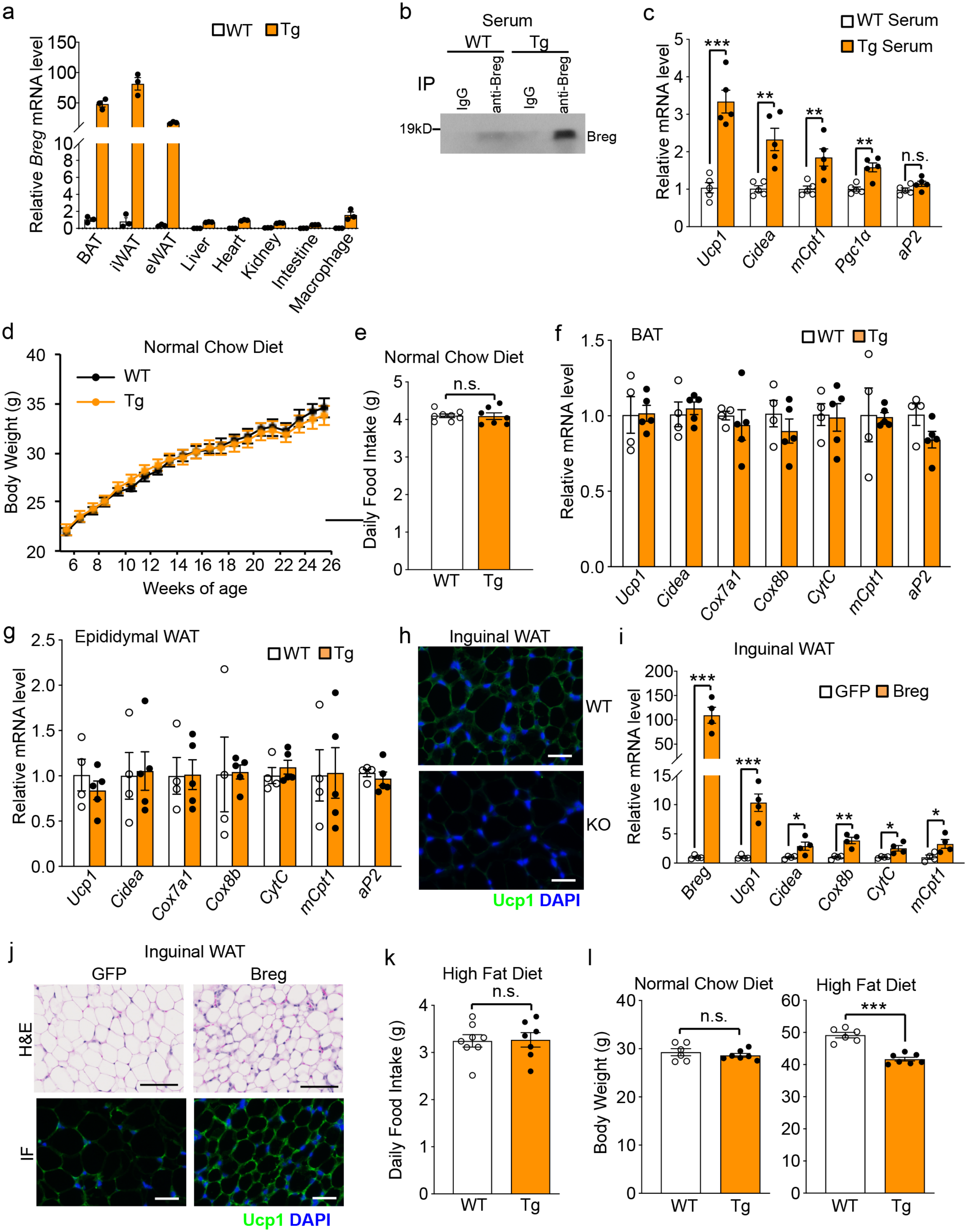

**Extended Data Fig. 3 | Phenotypes of *Breg* transgenic mice.** **a**, Relative mRNA levels of *Breg* in different tissues from *Breg* transgenic (Tg) mice and wild-type (WT) littermate controls (n=3 mice per group). **b**, Detection of Breg in WT and *Breg* Tg mice serum after immunoprecipitation with Breg antibody. **c**, Gene expression analysis in inguinal adipocytes treated with 10% serum from *Breg* Tg mice or WT controls (n=5). **d**, Body weight of *Breg* Tg mice (n=9) and WT controls (n=6) on normal chow diet. **e**, Daily normal chow food intake of 3-month-old male *Breg* Tg mice (n=7) and WT controls (n=8). **f**, **g**, Gene expression analysis in BAT (**f**) and epididymal WAT (**g**) from 2-month-old male *Breg* Tg mice (n=5) and littermate controls (n=4). **h**, Ucp1 immunofluorescence staining of inguinal WAT from WT and Ucp1 knockout mice. **i**, Gene expression analysis in inguinal WAT injected with *Breg* adenovirus (n=4 mice per group). **j**, H&E and Ucp1 immunofluorescence staining of inguinal WAT injected with *Breg* adenovirus (n=3 mice per group). **k**, Daily HFD food intake of *Breg* Tg mice (n=7) and littermate controls (n=8). **l**, Body weight of a second cohort of male *Breg* Tg mice (n=7) and littermate controls (n=6) before and after fed on 18-week of HFD. Scale bar, 200 μm. Data are mean ± s.e.m. *$p<0.05$, **$p<0.01$, ***$p<0.001$ and not significant (n.s.) by two-tailed Student's t test.

Extended Data Fig. 4

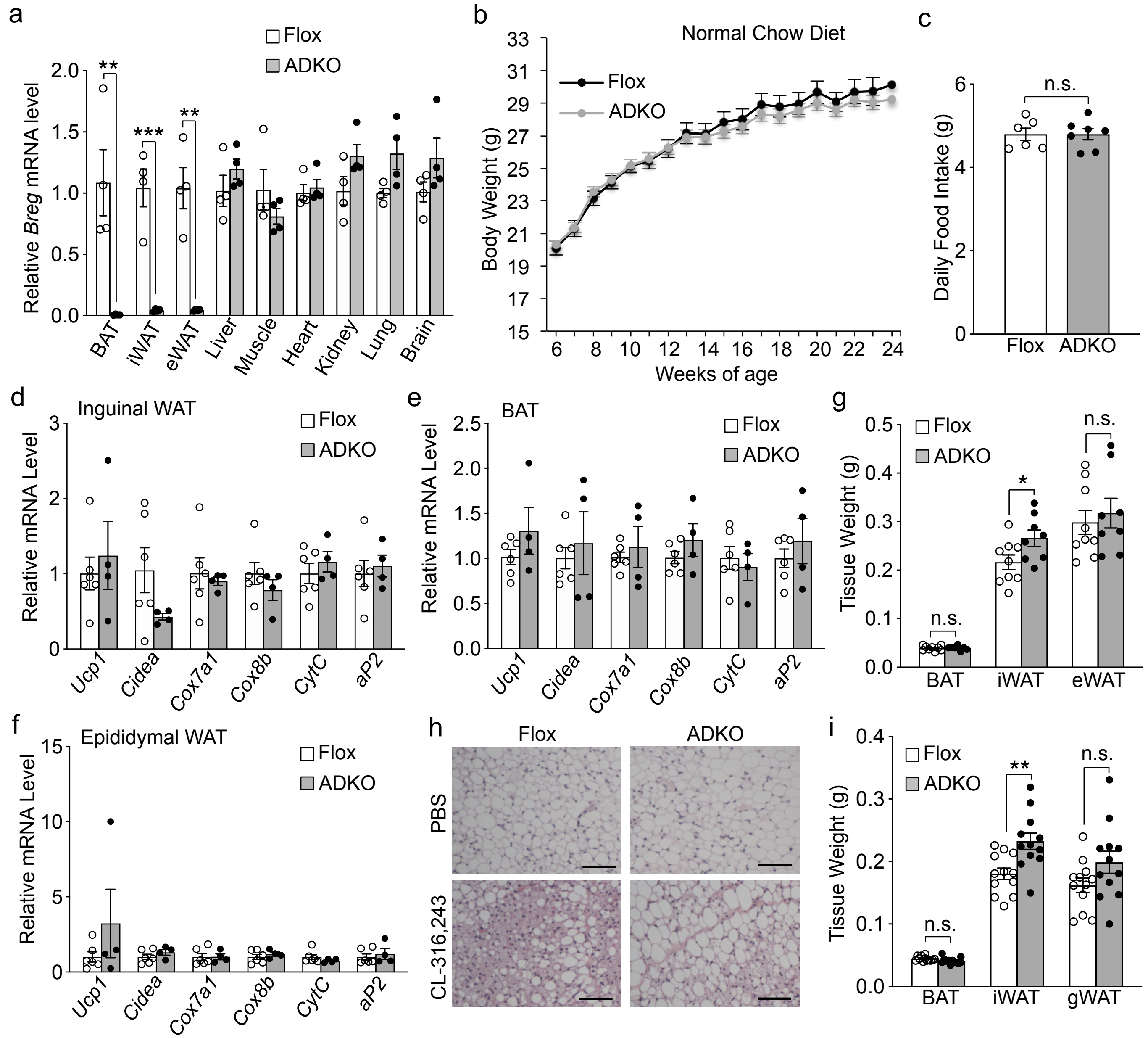


**Extended Data Fig. 4 | Phenotypes of *Breg* adipose specific knockout mice. a**, *Breg* expression in different tissues of *Breg* ADKO mice and Flox controls (n=4). **b**, Body weight of male *Breg* ADKO mice (n=11) and Flox controls (n=6) on normal chow diet. **c**, Daily normal chow food intake of male *Breg* ADKO mice (n=7) and Flox controls (n=6). **d-f**, Gene expression analysis of iWAT (**d**), BAT (**e**) and epididymal WAT (**f**) from 3-month-old male *Breg* ADKO mice (n=4) and Flox littermate controls (n=6) housed at 23 °C. **g**, Fat mass of 5-month-old female *Breg* ADKO mice (n=8) and Flox controls (n=9) after 7 hours cold exposure. **h**, Representative images of H&E staining of iWAT from 3-month-old female *Breg* ADKO mice and Flox controls after 2 days CL-316,243 administration (n=3 mice per group). **i**, Fat mass of mice from (**h**), (n=12). Scale bar, 200 μm. Data are mean ± s.e.m. *p<0.05, **p<0.01, ***p<0.001 and not significant (n.s.) by two-tailed Student's t test.

#### Extended Data Fig. 5

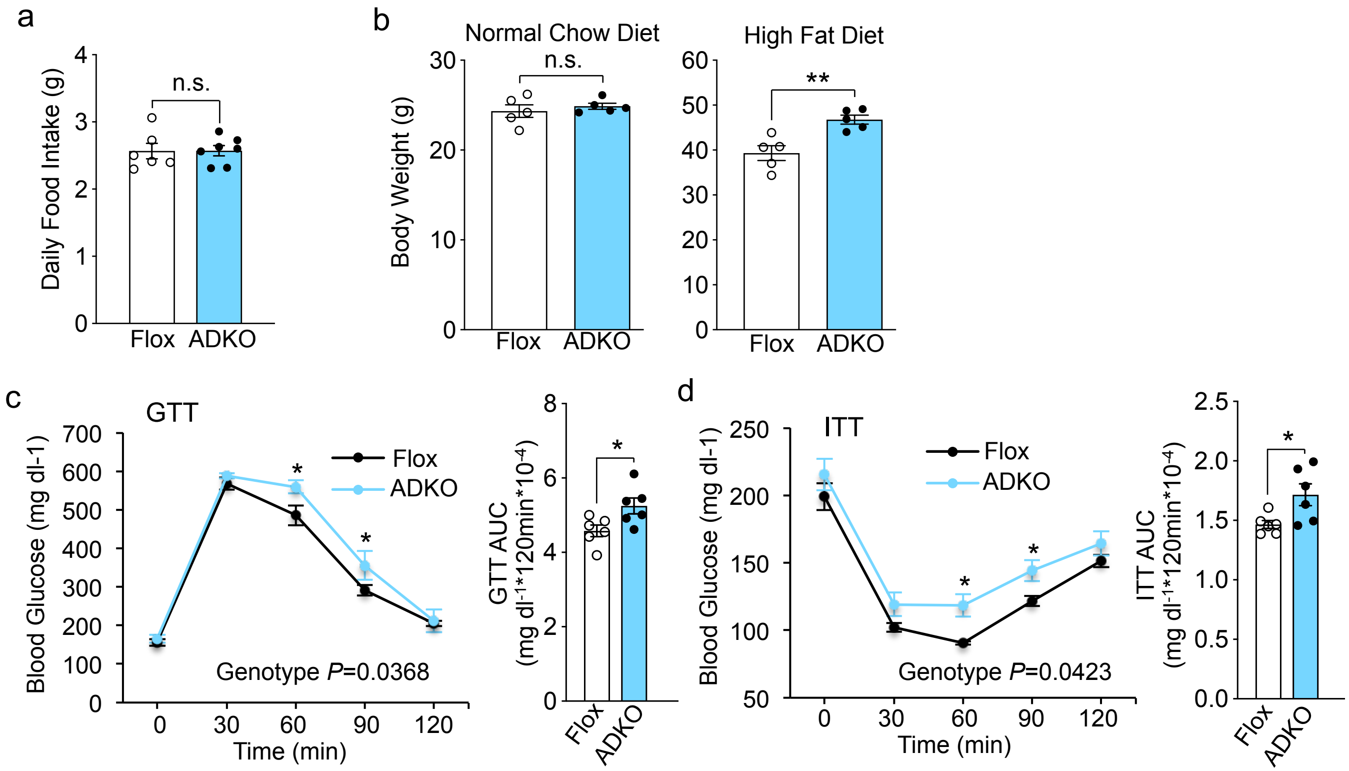


**Extended Data Fig. 5 | *Breg* ADKO mice had impaired glucose and insulin tolerance on HFD prior to body weight divergence from that of wild type mice. a**, Daily HFD food intake of male *Breg* ADKO mice (n=7) and Flox controls (n=6). **b**, Body weight of a second cohort of male *Breg* ADKO mice and Flox controls (n=5 mice per group) before and after fed on 18 weeks of HFD. **c**, GTT of *Breg* Flox and ADKO mice fed on 5 weeks of HFD (n=6 mice per group, Genotype p=0.0368). **d**, ITT of Breg Flox and ADKO mice fed on 6 weeks of HFD (n=6 mice per group, Genotype p=0.0423). Data are mean ± s.e.m. *p<0.05, **p<0.01, ***p<0.001 and not significant (n.s.) by two-tailed Student's t test (**a**, **b** and AUC in **c** and **d**) and two-way repeated measures ANOVA with post hoc test by Fisher's LSD test (**c** and **d**).

Extended Data Fig. 6

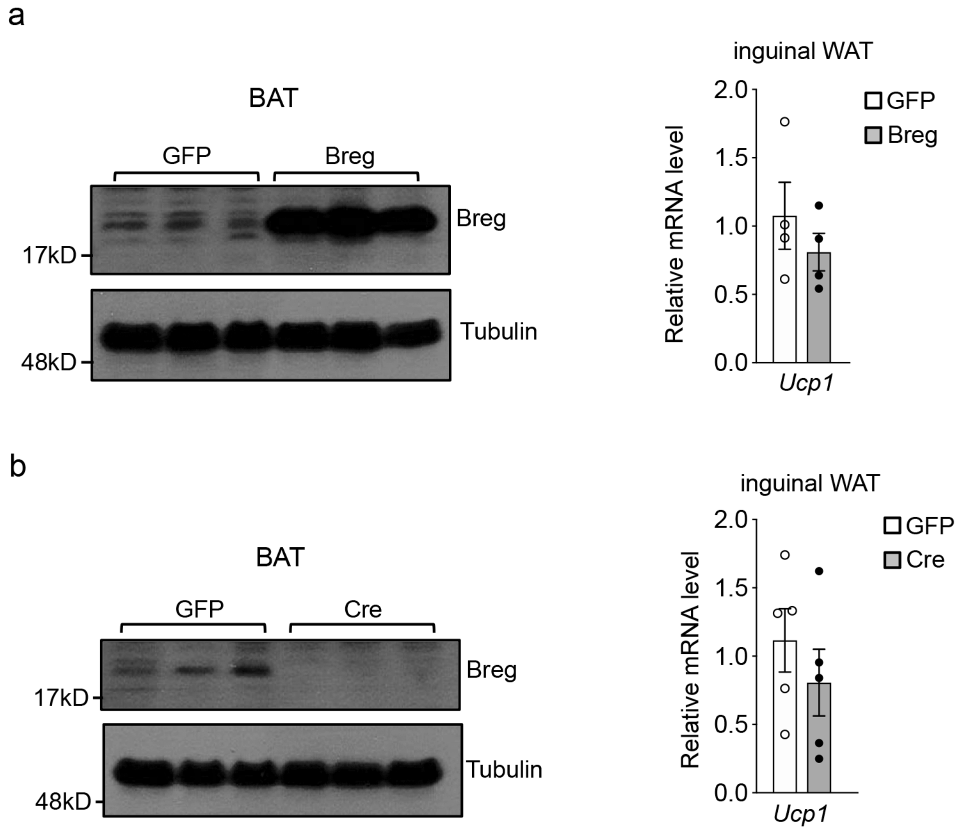


**Extended Data Fig. 6 | BAT-specific overexpression and deletion of *Breg* had no effect on *Ucp1* expression in iWAT. a**, Western blot of Breg in BAT and *Ucp1* expression in iWAT (n=4 mice per group) from BAT specific *Breg* overexpression mice. **b**, Western blot of Breg in BAT and *Ucp1* expression in iWAT (n=5 mice per group) from BAT specific *Breg* deletion mice. Data are mean ± s.e.m.

Extended Data Fig. 7

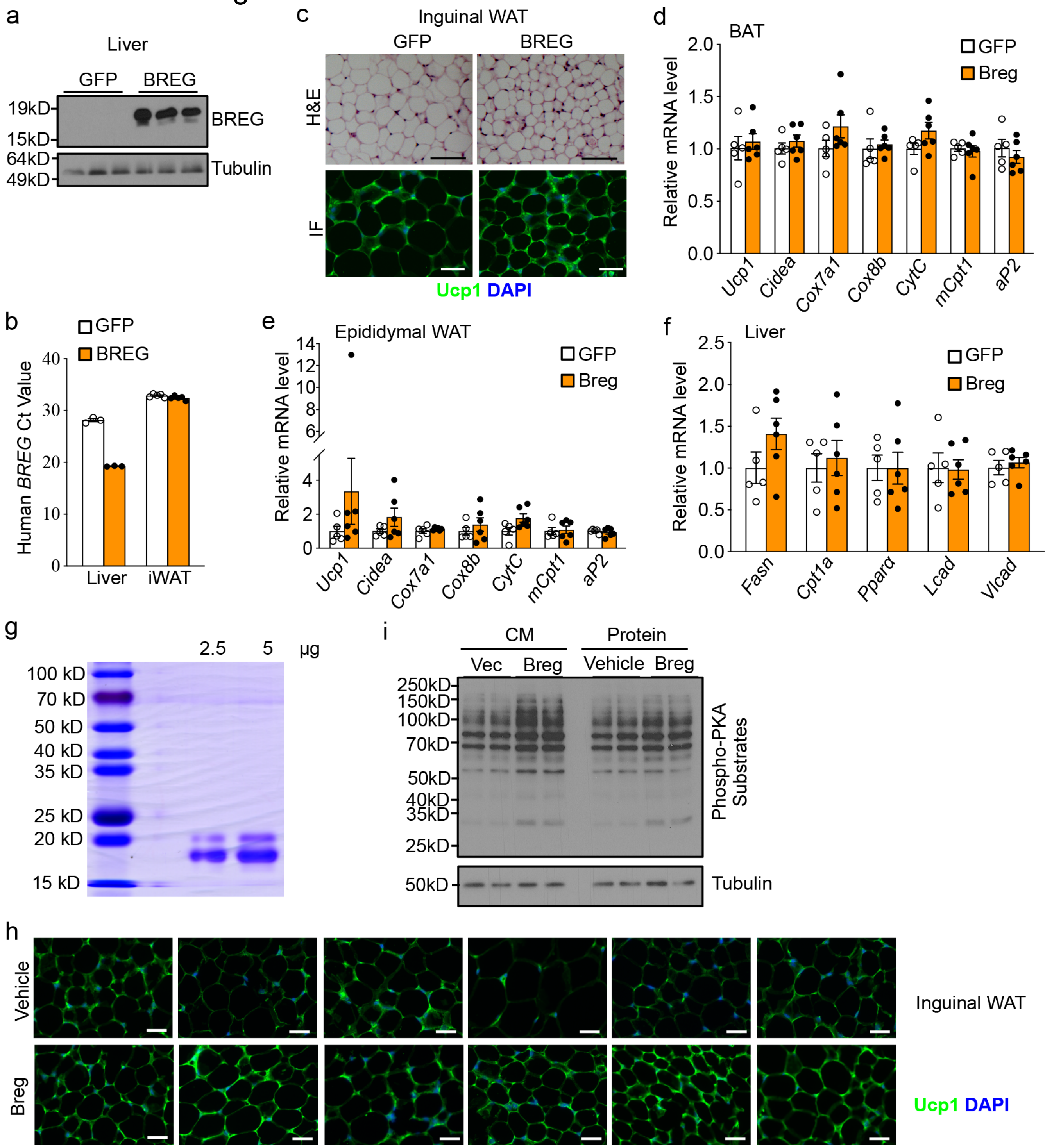


**Extended Data Fig. 7 | Circulating Breg induces WAT browning. a**, Western blot of BREG in liver from mice injected with *GFP* or human *BREG* adenoviruses. **b**, *BREG* RT-qPCR Ct value in liver and inguinal WAT from mice injected with *GFP* or *BREG* adenoviruses (n=3-5). **c**, Representative images of H&E staining and Ucp1 immunofluorescence staining of iWAT from mice injected with *GFP* or *BREG* adenoviruses (n=3 mice per group). Scale bar, 200 μm. **d-f**, Gene expression analysis in BAT(**d**), eWAT (**e**) and liver (**f**) from mice injected with *GFP* or *Breg* adenoviruses (GFP, n=5; Breg, n=6). **g**, SDS-PAGE gel of purified recombinant Breg protein. **h**, Ucp1 immunofluorescence staining of iWAT from mice injected with Breg protein or vehicle for 9 days. Scale bar, 200 μm. **i**, Western blot analysis of phosphorylated PKA substrates in brown adipocytes treated with conditioned medium (CM) containing 50 nM Breg protein or 50 nM purified Breg protein for 45 min. Data are mean ± s.e.m.

Extended Data Fig. 8

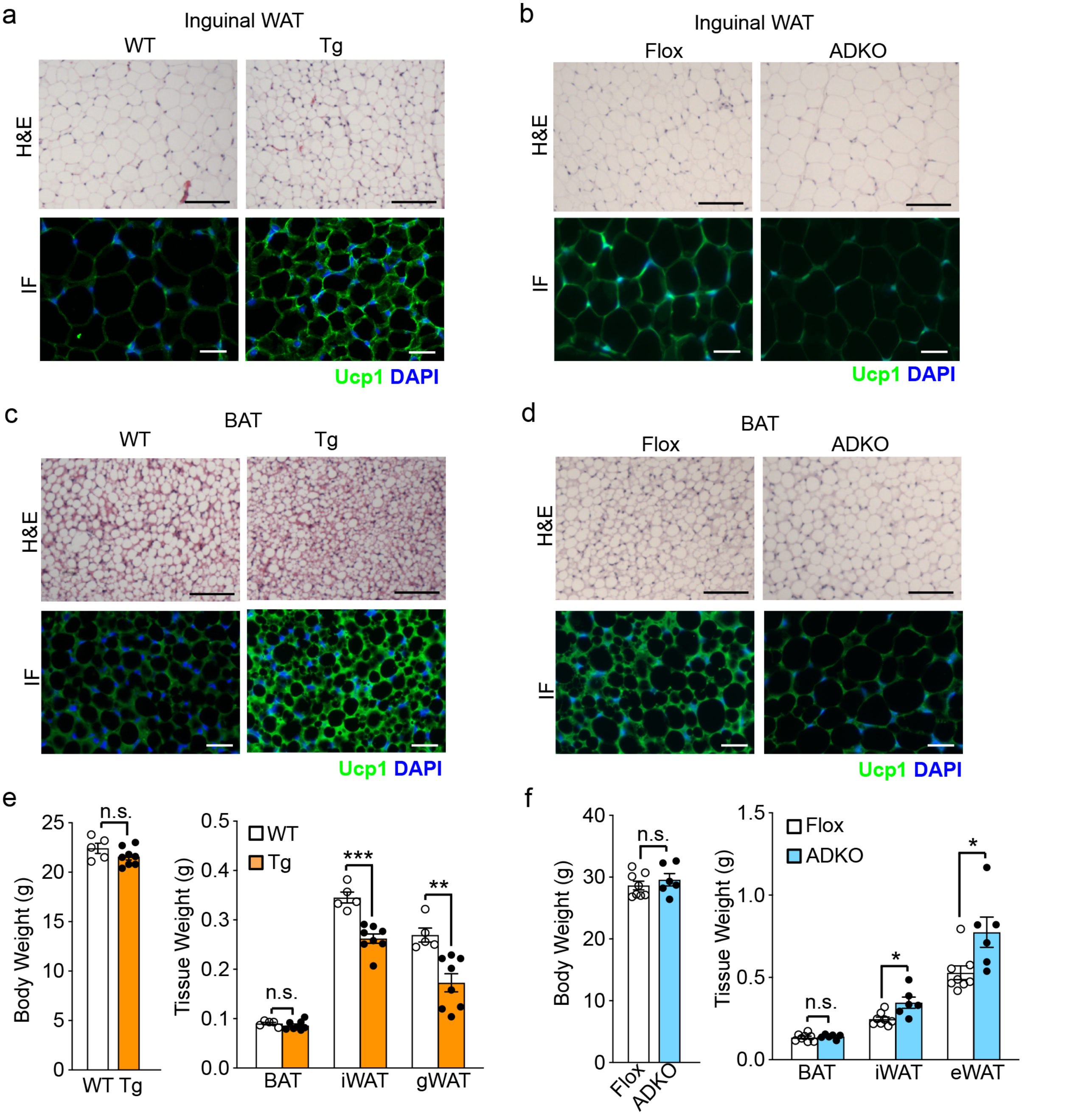


**Extended Data Fig. 8 | Breg functions independently of β-AR signaling. a**, **b**, H&E staining and Ucp1 immunofluorescence staining of inguinal WAT from *Breg* Tg female mice and WT controls housed at 30°C for 1 month (**a**) and from *Breg* ADKO male mice and Flox controls housed at 30°C for 2 months (**b**) (n=3 mice per group). **c**, **d**, H&E staining and Ucp1 immunofluorescence staining of interscapular BAT from *Breg* Tg female mice and WT controls housed at 30°C for 1 month (**c**) and from *Breg* ADKO male mice and Flox controls housed at 30°C for 2 months (**d**) (n=3 mice per group). **e**, Body weight and fat mass of *Breg* Tg female mice (n=8) and WT controls (n=5) housed at 30°C for 1 month. **f**, Body weight and fat mass of *Breg* ADKO male mice (n=6) and Flox controls (n=8) housed at 30°C for 2 months. Scale bar, 200 μm. Data are mean ± s.e.m. *p<0.05, **p<0.01, ***p<0.001 and not significant (n.s.) by two-tailed Student's t test.

Extended Data Fig. 9

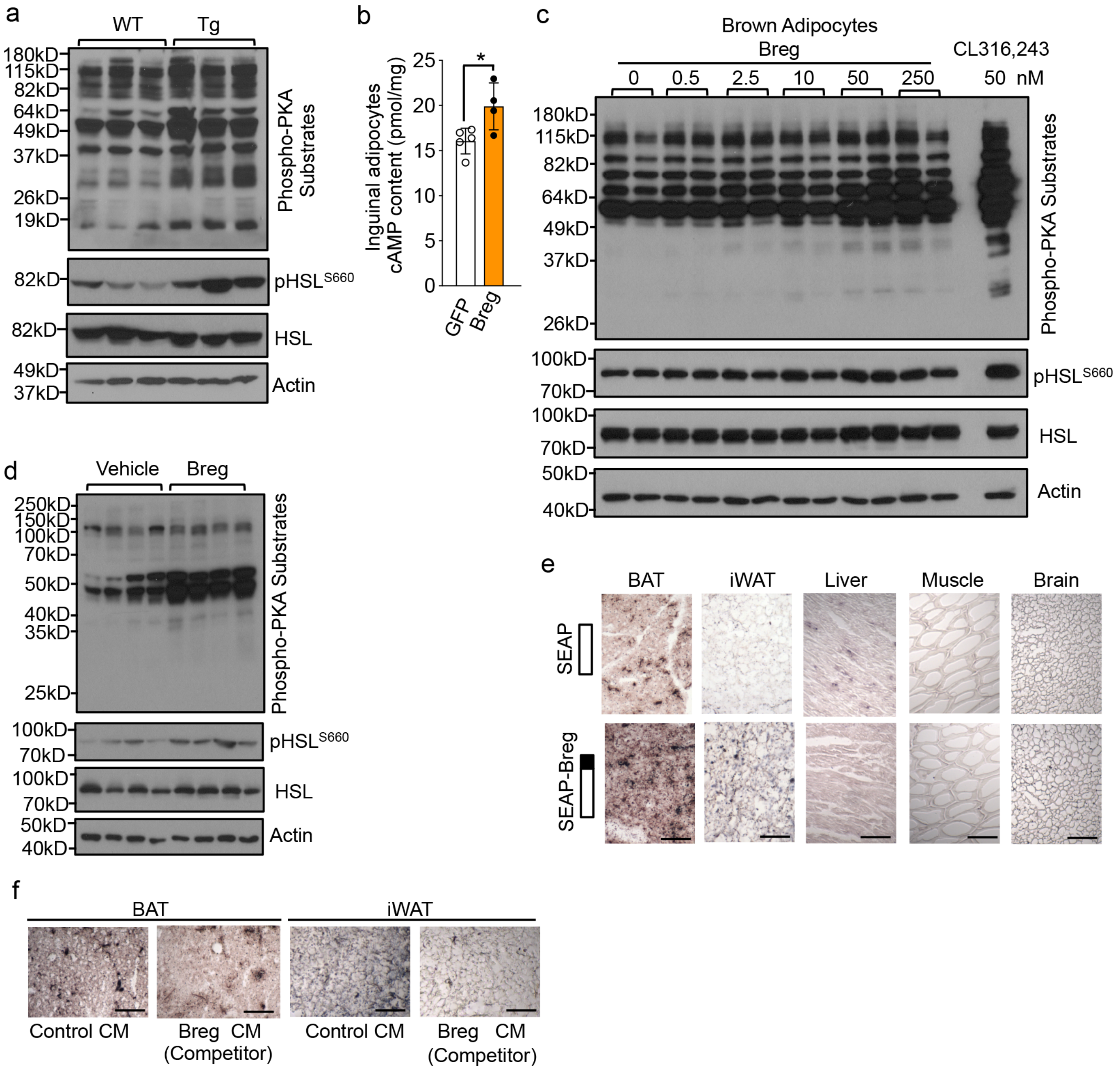


**Extended Data Fig. 9 | Breg functions though PKA signaling pathway. a**, Western blot analysis of phosphorylated PKA substrates, phospho- and total HSL in iWAT from *Breg* Tg mice and littermate controls housed at 23°C. **b**, cAMP levels in primary inguinal adipocytes transduced with *Breg* (n=4) or *GFP* (n=5) adenoviruses. **c,** Western blot analysis of phosphorylated PKA substrates, phospho- and total HSL in brown adipocytes treated with different dose of Breg protein for 30 min. **d**, Western blot analysis of phosphorylated PKA substrates, phospho- and total HSL in iWAT from 3-month-old WT male mice after 9 days of Breg administration. **e**, SEAP or SEAP-Breg binding on indicated frozen tissue sections. Scale bar, 200 μm. **f**, SEAP-Breg binding to BAT and iWAT frozen sections in the presence of indicated conditioned medium (CM). Scale bar, 200 μm. Data are mean ± s.e.m. *$p<0.05$ by two-tailed Student's t test.
